# Ubiquitous bacterial polyketides induce cross-kingdom microbial interactions

**DOI:** 10.1101/2022.05.09.491136

**Authors:** Mario K. C. Krespach, Maria C. Stroe, Tina Netzker, Maira Rosin, Lukas M. Zehner, Anna J. Komor, Johanna M. Beilmann, Thomas Krüger, Olaf Kniemeyer, Volker Schroeckh, Christian Hertweck, Axel A. Brakhage

## Abstract

Although the interaction of prokaryotic and eukaryotic microorganisms is critical for the functioning of ecosystems, knowledge of the processes driving microbial interactions within communities is in its infancy. We previously reported that the soil bacterium *Streptomyces iranensis* specifically triggers the production of natural products in the fungus *Aspergillus nidulans.* Here, we discovered that arginine-derived polyketides serve as the bacterial signals for this induction. Arginine-derived polyketide-producing bacteria occur world wide. These producer bacteria and the fungi that decode and respond to this signal can be co-isolated from the same soil sample. Arginine-derived polyketides impact surrounding microorganisms both directly as well as indirectly, by inducing the production of natural products in fungi that further influence the composition of microbial consortia.

**One-Sentence Summary:** Ubiquitous bacterial polyketides are universal components of the chemical network for microbial communication

## Main Text

In all known habitats on earth microorganisms form consortia with a multitude of prokaryotic and eukaryotic microorganisms (*1*). Nearly every day new data concerning the composition of microbial consortia are published. They impressively demonstrate the diversity of microorganisms in various ecosystems (*2*) that provide services critical for life. For example, soil filters and stores water, provides a medium that supplies plants and heterotrophic organisms with water and nutrients, offers a habitat for a large diversity of organisms, and is the source of most of our antibiotics. Many of the vital soil functions are due to the activity of microorganisms that regulate nutrient cycling, decompose organic matter, define soil structure, suppress plant diseases, and support plant productivity (*3, 4*). However, despite the importance of microbial consortia for a healthy ecosystem, the elucidation of functional interactions between bacteria and fungi that determine the composition of microbial consortia is still in its infancy. These interactions are decisive for the functioning of microbial communities. One example is lichens, microbial communites composed of fungi and phototrophic microorganisms like algae or cyanobacteria (*5*) that are able to colonize extreme environments like rocks and fix carbon dioxide even under harsh conditions (*6*). Lichens also provide microhabitats for many bacteria thus forming a complex microbial consortium (*7*). Likewise, it was shown that microorganisms from different kingdoms drive the assembly of microbiota in preterm infants (*8*).

We are beginning to understand that microorganisms communicate with and influence each other *via* a chemical language composed of low molecular weight organic compounds that are part of the greater chemical category commonly referred to as natural products (NPs) (*9, 10*). Some of these compounds have been assigned functions with reference to their impact on humans, *i.e*. as antibiotics or toxins (*1, 11, 12*). However, the ecological role for most of these compounds remains obscure. What has been shown is that the ecological context, including the presence of other microorganisms, is important for production of these compounds (*13–15*). Consequently, many of the gene clusters in microorganisms encoding the biosynthesis of such NPs are silent under conventional laboratory conditions (*12*) and biosynthesis can only be activated when the correct microbial partner is provided. We reported an early example of this scenario when we showed that the bacterium *Streptomyces rapamycinicus* and its closest relative *Streptomyces iranensis* specifically trigger the expression of the otherwise silent orsellininc acid (*ors*) gene cluster of the fungus *Aspergillus nidulans,* which results in the production of orsellinic acid and derivatives thereof (*14, 16*). However, the bacterial agent triggering the fungal production of NPs remained elusive. Here, we report the discovery of one such class of molecules in the ubiquitous bacterial arginine-derived polyketides that both directly impact surrounding microorganisms and specificaly induce the production of NPs in fungi that further influence the composition of microbial consortia.

## Results

### A functional bld regulon in S. iranensis is essential for the induction of the fungal ors gene cluster

In order to identify the bacterial trigger of the silent fungal *ors* biosynthetic gene cluster (BGC), we analyzed the well-characterized pleiotropic Bld regulators of streptomycetes (Figure 1A). The designation of these genes as *bld* (for bald) originates from the growth phenotype shown by the corresponding mutants, *i.e.,* they lack formation of aerial mycelium and do not develop the typical fuzzy appearance of *S. iranensis* colonies, but instead stay smooth (*17*). Because BldD is at the top of the gene regulation cascade involved in the developmental cycle of streptomycetes as well as in natural product biosynthesis (*18*) (Figure 1A), we deleted its encoding gene in *S. iranensis*. The generated mutant displayed the “bald-colony” phenotype and, as shown by LC-MS analysis, co-cultures of the mutant and *A. nidulans* did not contain *ors* BGC-derived orsellinic acid (Figure 1A, B). To further substantiate our finding, we also deleted two other genes of the BldD regulon in *S. iranensis*, *bldH* that encodes a pleiotropic regulator of development and whose transcription is controlled by BldD (*19*) and *bldA* encoding the only tRNA in most streptomycetes that is able to effectively translate the TTA codon into leucine (*20, 21*) (Figure 1A). Likewise, the generated *bldA* and *bldH* deletion mutants of *S. iranensis* exhibited the expected bald phenotype and did not induce the appearance of orsellinic acid in the supernatant of co-cultures with *A. nidulans* (Figure 1A, B). This data indicated that genes controlled by the BldD regulon regulate the bacteria-induced activation of the fungal *ors* BGC.

**Fig. 1:**
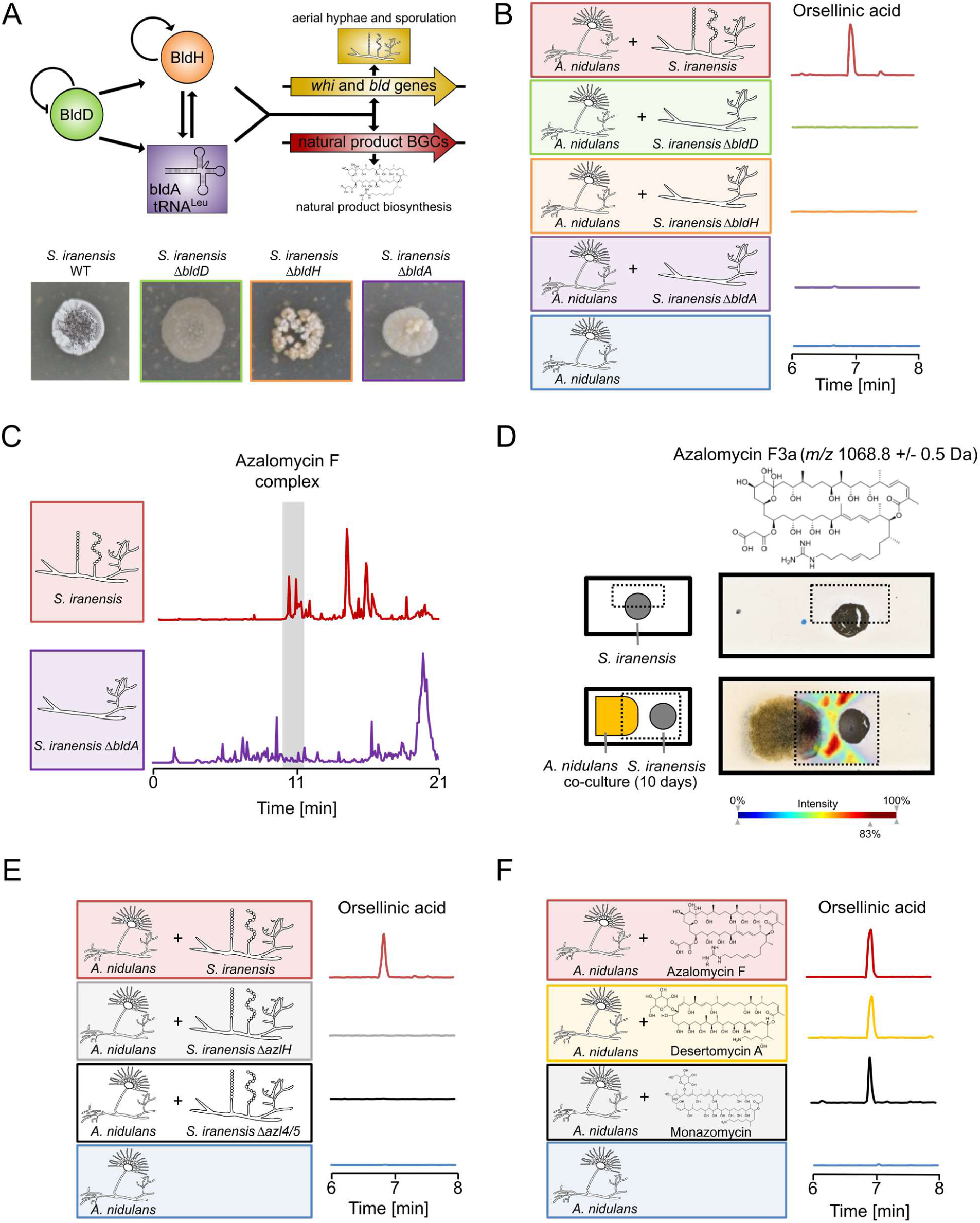
Identification of *Streptomyces*-derived marginolactones as inducing agents for production of orsellinic acid and derivatives by the fungus *A. nidulans*. **A:** Top: Regulatory cascade of *bld* genes. Bottom: Deletion mutants of *bldD*, *bldH*, and *bldA* in *S. iranensis* lack formation of aerial hyphae and spores. **B:** Co-culture of *A. nidulans* with *S. iranensis* WT and mutant strains Δ*bldD,* Δ*bldH,* Δ*bldA* and extracted ion chromatogram for orsellinic acid (*m/z* 167 [M–H]^-^) derived from LC-MS analysis of culture supernatant. **C:** Total ion chromatogram of culture extracts of *S. iranensis* wild type and Δ*bldA.* Missing peaks correspond to the azalomycin F complex only found in the *S. iranensis* wild-type strain. **D:** MALDI-IMS analysis of co-cultivation of *S. iranensis* and *A. nidulans* for 10 days on agar. Left: Schematic visualization of sample preparation. *S. iranensis* is portrayed in grey, *A. nidulans* in yellow. The box indicates the measured area. Right: MALDI-IMS analysis of the distribution of azalomycin F3a (*m/z* 1068.8 ± 0.5 Da). Abundance of the analyzed mass is depicted as a heat map from low abundance (blue) to high abundance (red). **E:** Co-cultivation of *A. nidulans* with azalomycin F-deficient *S. iranensis* mutants and extracted ion chromatogram of orsellinic acid (*m/z* = 167 [M-H]^-^) derived from LC-MS analysis of the co- culture extract. **F:** Cultivation of *A. nidulans* with indicated marginolactones and extracted ion chromatogram of orsellinic acid (*m/z* 167 [M-H]^-^) derived from LC-MS analysis of the culture extract.

### Bld-regulon regulates mRNA, protein and product level of azalomycin F biosynthesis in S. iranensis

In order to identify genes and compounds regulated by the *bld*-regulon, we performed transcriptome, proteome and LC-MS-based metabolome analyses of the *bldA* deletion mutant compared with the wild-type strain. We selected the *bldA* mutant, since its regulatory power is well documented. It directly controls all genes carrying a TTA codon and hence target genes are easy to identify. In the Δ*bldA* mutant strain, downregulation of many genes and proteins, including several genes and proteins of the azalomycin F BGC, was observed (Table 1). This finding was reflected by the lack of molecular masses corresponding to azalomycin F in culture extracts of the Δ*bldA* mutant strain compared to wild type (Figure 1C).

**Table 1:**
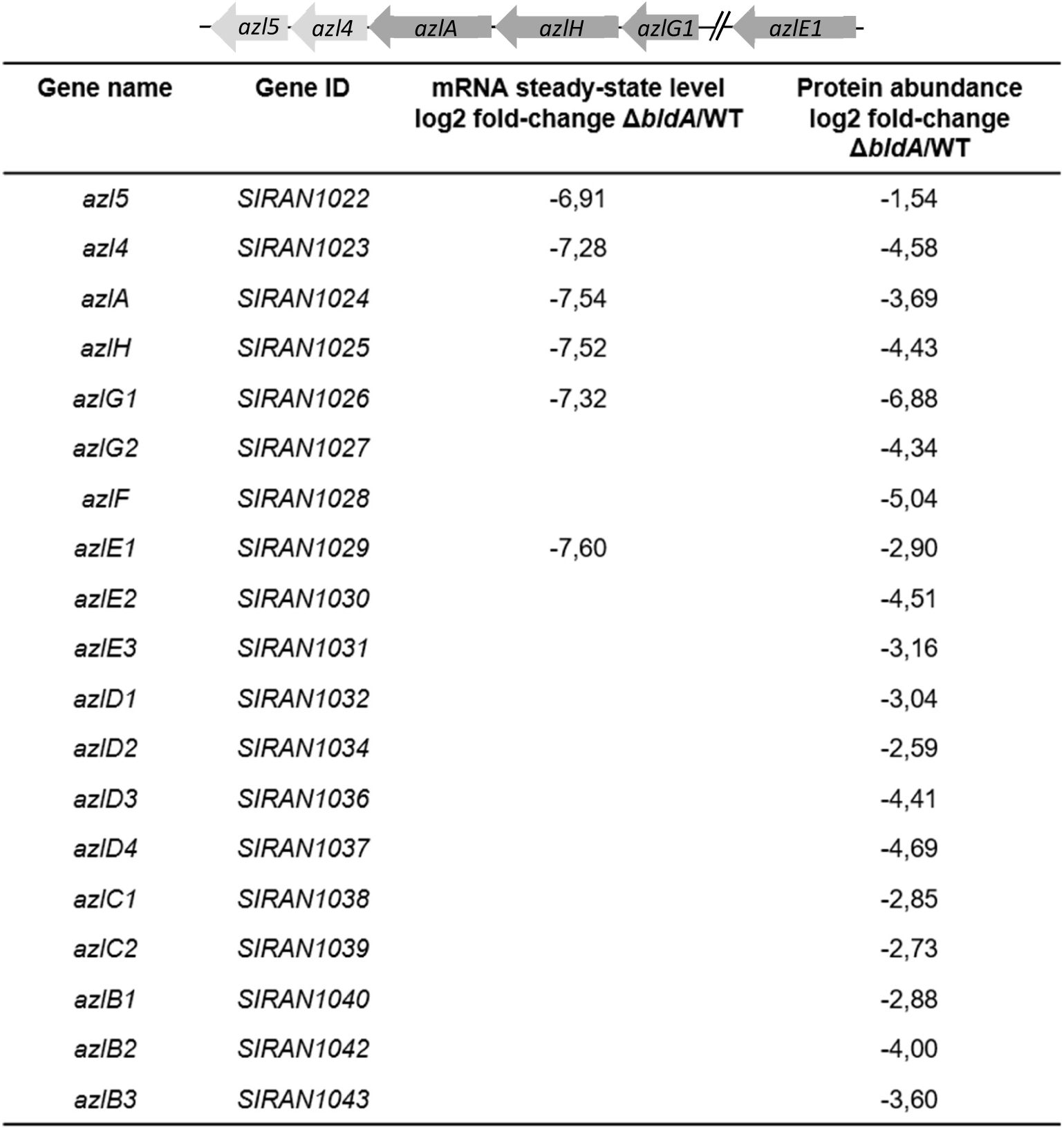
Changes in gene expression and relative protein abundance in the *S. iranensis* Δ*bldA* mutant compared to wild type. mRNA steady-state level and relative protein abundance of core biosynthetic genes/ proteins (depicted on top) of the azalomycin F BGC. Data are depicted as log2 fold-changes of TPM (Transcripts Per Kilobase Million) values and protein abundance measured in the Δ*bldA* mutant compared to the wild type.

### Azalomycin F is overproduced and secreted by S. iranensis in presence of A. nidulans

To analyze and visualize a potential release of azalomycin F towards the fungus, we applied matrix-assisted laser desorption-ionization imaging mass spectrometry (MALDI-IMS) to co-cultures of *S. iranensis* and *A. nidulans*. Two metabolites likely representing the azalomycin F complex were detected in the confrontation zone, consistent with azalomycin F3a (*m/z* 1069.12) and azalomycin F4a (*m/z* 1083.16) (*22*) (Figure 1D, Supplementary Figure 1). These compounds accumulated at the side of the bacterial colony facing *A. nidulans*, suggesting that the release of azalomycin F is directed towards the fungus. After ten days of co-cultivation (Figure 1D, Supplementary Figure 1), azalomycin F co-localized with the fungus. This finding agrees with our LC-MS data indicating that azalomycin F accumulates in the biomass fraction of an *A. nidulans* monoculture supplemented with azalomycin F (Supplementary Figure 2).

### Azalomycin F triggers NP biosynthesis in A. nidulans

To demonstrate that azalomycin F is a bacterial signal triggering fungal NP biosynthesis, we analyzed mutant strains with deletion of azalomycin F biosynthesis genes (*16, 23, 24*). Co-cultivation of the respective *S. iranensis* mutant strains Δ*azlH* and Δ*azl4*Δ*azl5* with *A. nidulans* did not result in the production of orsellinic acid and its derivatives (Figure 1E; Supplementary Figure 3A). As an important proof, we added purified azalomycin F to monocultures of *A. nidulans*. It clearly induced the production of orsellinic acid, lecanoric acid, and derivatives F-9775A and B (Figure 1F, Supplementary Figure 4A). The production of compounds was accompanied by increased mRNA steady-state levels of the orsellinic acid biosynthesis gene *orsA* (Supplementary Figure 3B). Collectively, azalomycin F is the sought-after bacterial trigger that activates NP biosynthesis in *A. nidulans*.

### Azalomycin F and other marginolactones specifically induce fungal NP biosynthesis

Azalomycin F is a member of the family of marginolactones whose biosynthesis starts with arginine. Marginolactones consist of a macrolactone backbone and a side chain containing a guanidyl- or amino group (*25*). The marginolactones desertomycin A and monazomycin are produced by *Streptomyces macronensis* and *Streptomyces mashuensis*, respectively (*26, 27*). Since we found that purified desertomycin A and monazomycin also induced the production of orsellinic acid and its derivatives in *A. nidulans* (Figure 1F, Supplementary Figure 4A), we proposed that actinomycetes producing these compounds are also able to induce the fungal *ors* gene cluster. This is indeed the case, as co-cultures of *S. macronensis* and *S. mashuensis* with *A. nidulans* contained orsellinic acid and derivatives (Supplementary Figure 5A).

We also tested oasomycin B (*28*), a desertomycin-family compound lacking the amino group in its side chain. The importance of the amino moiety is reflected by the observation that oasomycin B had lost the antibacterial activity assigned to desertomycin A and that the reconstitution of a positively charged moiety to oasomycin restores antibacterial activity (*29*). In comparison to desertomycin A, equimolar concentrations of oasomycin B minimally triggered the production of orsellininc acid and its derivatives by the fungus (Supplementary Figure 4A). Thus, we concluded that the positively charged moieties in marginolactones are required for the induction of fungal NP biosyntheses. To determine the specificity of the inducing activity of marginolactones, we tested known antifungal compounds which either act on the cell wall biosynthesis, like caspofungin, or on the fungal membrane, such as amphotericin B or voriconazole (*30*). In contrast to marginolactones, none of the structurally different antimycotics induces the *ors* BGC (Supplementary Figure 4B).

### Worldwide distribution of arginine-derived polyketide-producing bacteria

Our findings suggested that bacteria capable of producing marginolactones are inducers of cryptic BGCs in fungi. To facilitate and accelerate testing of compounds and strains for their ability to induce the *ors* BGC, we generated an *A. nidulans* strain that had the promoter of the orsellinic acid biosynthesis gene *orsA* (*orsA*p) fused with a translational fusion of the gene encoding a nanoluciferase (nLuc) with the green fluorescence gene (GFPspark, GFPs). The gene fusion was integrated at the *orsA* locus in the genome of *A. nidulans* and is thus controlled by the *orsA* promoter (Figure 2A; Supplementary Figure 6). In this reporter strain, induction of the *ors* gene cluster was readily detectable by GFPs-dependent green fluorescence of mycelia and could be quantified via the activity of nLuc.

**Fig. 2:**
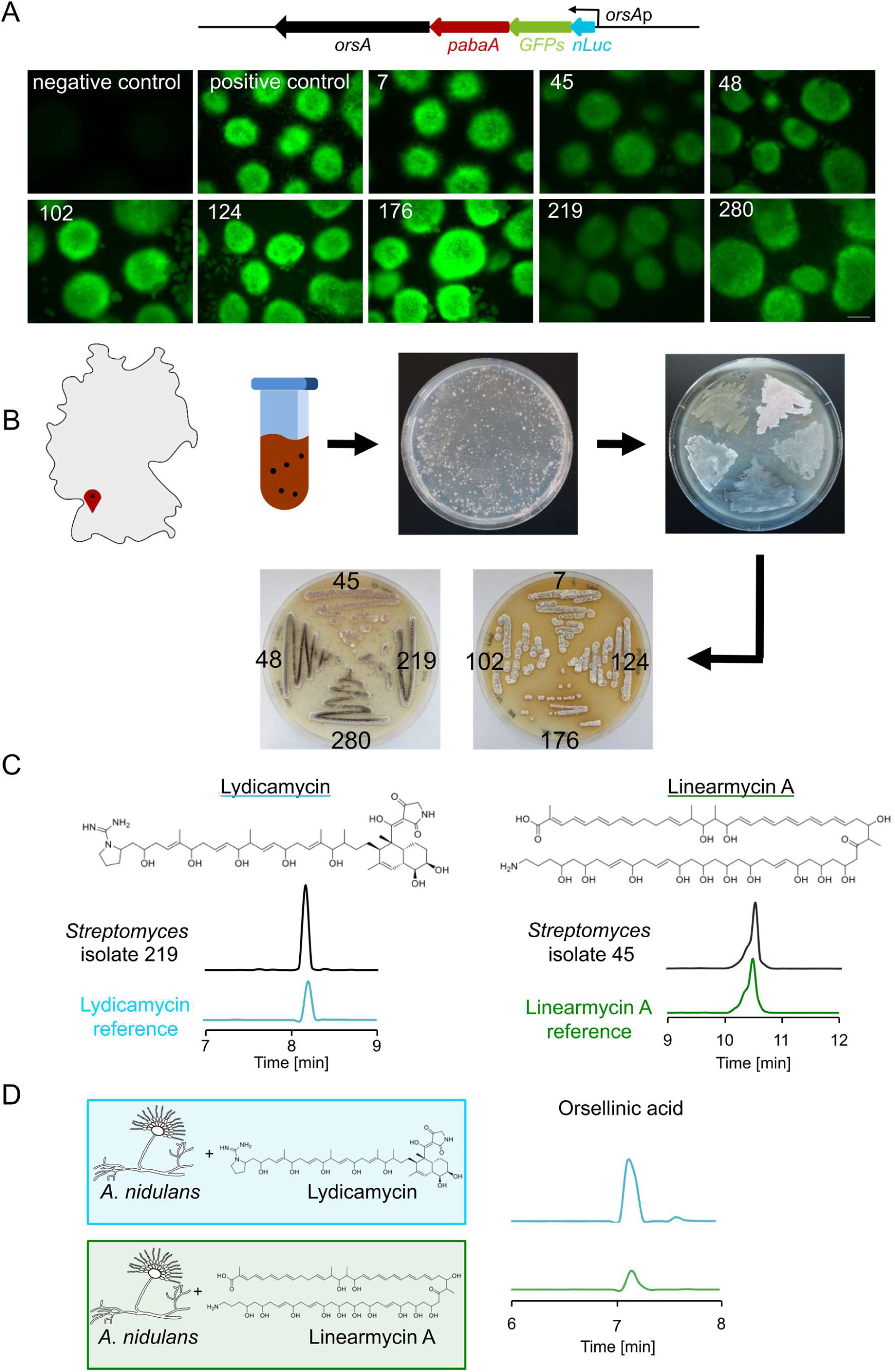
Isolation of bacteria from soil producing arginine-derived polyketides and inducing the *ors* BGC *of A. nidulans*. **A:** On top schematic depiction of the *orsA*p-nLuc-GFPs reporter gene fusion. Pictures show *A. nidulans orsA*p-nLuc-GFPs reporter strain co-cultured for 6 h with *Streptomyces* isolates 7, 45, 102, 124, 176, 219 an 280 isolated from soil. Negative control: *A. nidulans orsA*p-nLuc-GFPs without addition of bacteria. Positive control: *A. nidulans orsA*p-nLuc-GFPs co-cultured with known inducer *S. iranensis*. Scale bar: 500 µm. **B:** Map of Germany with marked location of the origin of soil sample and workflow for isolation of bacteria from soil. **C:** Cultivation of soil isolates *Streptomyces* sp. 219 and 45 and extracted ion chromatogram for lydicamycin (*m/z* 841 [M+H]^+^)) and linearmycin A (*m/z* 1140 [M+H]^+^) derived from LC-MS analysis of the culture supernatant. Structure of lydicamycin and linearmycin A are shown on top. **D**: Cultivation of *A. nidulans* with either lydicamycin or linearmycin and extracted ion chromatogram of orsellinic acid derived from LC-MS analysis of the culture extract.

Next, to determine the frequency of occurrence of marginolactone producers in nature, we screened soil for their presence. From 600 mg of soil we could cultivate 305 filamentously growing bacteria representing potential actinomycetes (Figure 2B). When the isolated strains were co-cultured with the *A. nidulans orsA*p*-nLuc-GFPs* reporter strain, eight of them triggered fluorescence (Figure 2A). The genomes of these bacterial strains were sequenced by Illumina sequencing. Using AntiSMASH and BlastN the obtained genomes were analyzed for the unusual arginine-loading domain in polyketide synthases required for biosynthesis of arginine-derived polyketides. All eight strains contained BGCs for the biosynthesis of arginine-derived polyketides (Supplementary Tables 1 and 2). The genomes of bacterial isolates 7, 48, 102, 124, 176, 219, and 280 bear a potential lydicamycin BGC, and isolate 45 a BGC for linearmycin A. This finding is interesting because it extends the spectrum of inducing marginolactones to include linear arginine-derived polyketides. Subsequent HPLC-MS analysis of the culture extracts from bacterial isolates 219 and 45 comfirmed the production of lydicamycin and linearmycin (Figure 2C). Further, both compounds induced the formation of orsellinic acid and its derivatives when added to cultures of *A. nidulans* (Figure 2D). We hypothesized that the producing strains should also be capable of inducing the fungal compounds. This was the case, as formation of orsellinic acid and derivatives was triggered by the *Streptomyces* isolates 219 and 45 producing lydicamycin and linearmycin, respectively (Supplementary Figure 8B). Therefore, not only marginolactones, but also linear arginine-derived polyketides, are able to induce fungal NP biosynthesis.

To search for arginine-derived polyketide producers worldwide, we followed two approaches: We screened (i) published data for regions where producers of azalomycin F, desertomycin A, monazomycin, linearmycin A, and lydicamycin were found and (ii) available genome sequences for presence of genes encoding the arginine-loading domain required for azalomycin F biosynthesis (Figure 4A). Numerous actinomycetes meet one or both of the criteria. These actinomycetes had been sampled all over the world on virtually all continents, which underlines their ubiquitous distribution (Figure 4A, Supplementary Table 3). Thus, it is very likely that there are far more species and strains producing the different arginine-derived polyketides.

It is challenging to identify NPs in the soil using analytical techniques because of their tendency to adsorb to soil particles (*31*). We approached this problem with our sensitive *A. nidulans* reporter strain, which fluoreces in response to induction of the *ors* cluster via *orsAp- nLuc-GFPs* expression, and therefore indicates the presence of an inducer molecule.Addition of supernatant of soil to a culture of the *A. niduans* reporter strain led to a clear increase in activity of the nLuc compared to a culture without soil supernatant (Supplementary Figure 12). Because the *ors* cluster is specifically induced by the arginine-derived polyketide marginolactone, this data suggests that arginine-derived polyketides are indeed present in soil.

### Frequency of occurrence of fungal responders to the arginine-derived polyketide signal

To obtain insights into whether arginine-derived polyketides are processed by fungi other than *A. nidulans,* we analyzed *Aspergillus fumigatus*, a human pathogen causing life-threatening infections. Despite being phylogenetically distantly related to *A. nidulans* (https://www.ncbi.nlm.nih.gov/genome/?term=Aspergillus%20nidulans), *A. fumigatus*

(https://www.ncbi.nlm.nih.gov/genome/?term=aspergillus+fumigatus) also specifically reacts to *S. rapamycinicus* and *S. iranensis* by the activation of the BGCs for fumicyclines and fumigermin (*13, 32*). When we added purified azalomycin F to monocultures of *A. fumigatus* the fumicyclines as well as fumigermin were detected in the culture, indicating that also this fungus responds to azalomycin F (Supplementary Figure 7).

Motivated by this finding, we sought to determine the frequency of occurrence of potential signal-decoding fungi. For this purpose, we isolated filamentous fungi from the same soil sample as used for the isolation of filamentous bacteria (Figure 3A). In total, we isolated 106 fungal strains and tested their repsonse to the *S. iranensis* WT and the corresponding *S. iranensis* Δ*azlH* mutant strain. Of the 106 fungal strains, 31 showed a change in culture coloration when co-cultured with *S. iranensis* WT that was not seen for the Δ*azlH* deletion mutant, suggesting induced production of NPs (Supplementary Figure 9A). For one strain we were able to identify the fungal compound whose production was triggered by *S. iranensis*. The analysis of HPLC-MS spectra indicated that fungal isolate 27 produces carviolin only in presence of *S. iranensis* WT but not Δ*azlH* (Figure 3B). Carviolin is a red pigment of *Penicillium* spp. (*33*), with potential silkworm-attracting properties (*34*). In agreement with this finding, fungal isolate 27 overproduced carviolin also in the presence of the linearmycin-producing bacterial isolate 45 and the lydicamycin-producing bacterial isolate 219 (Supplementary Figure 9B).

**Fig. 3:**
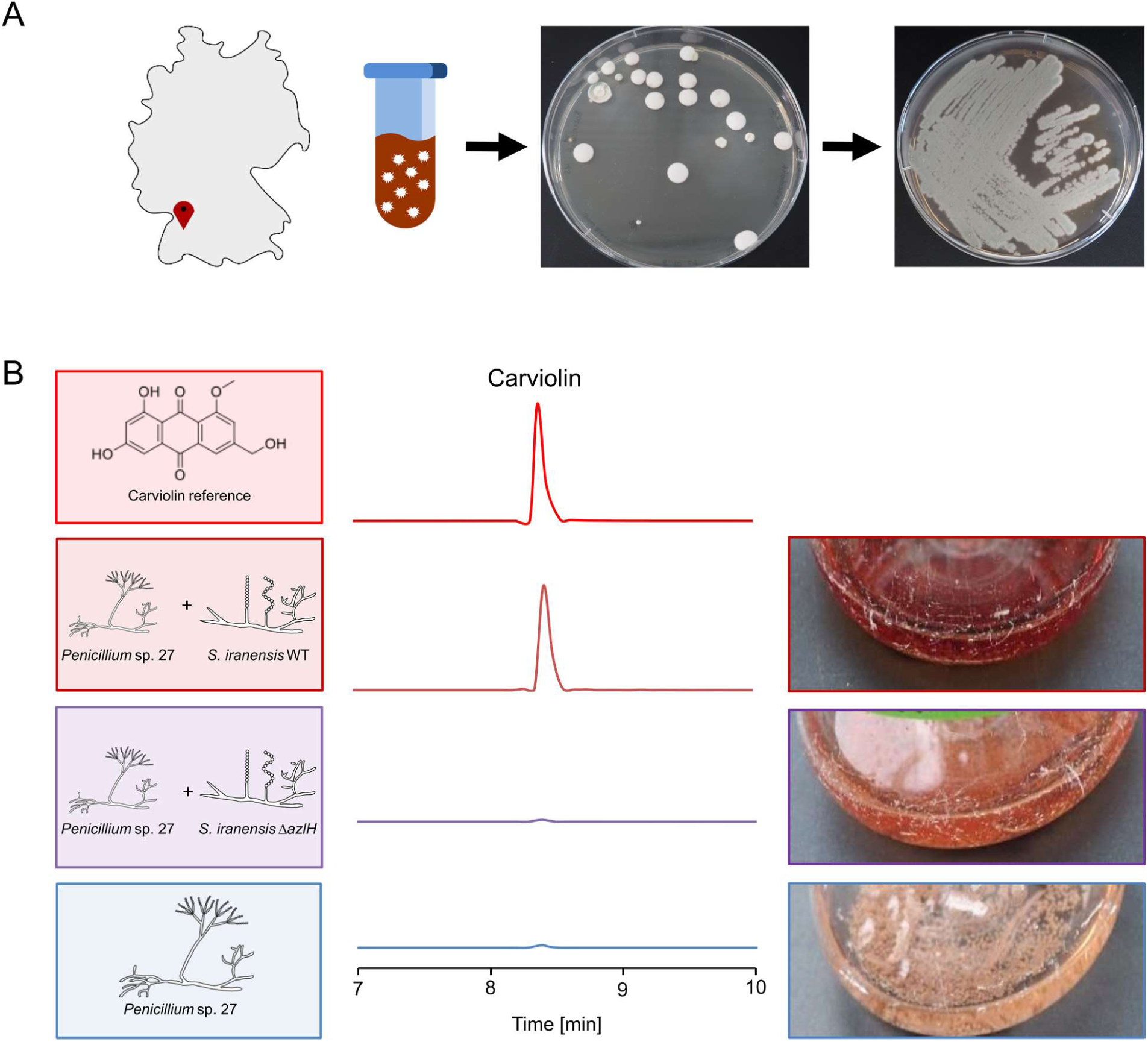
Isolation of *Penicillium* isolate 27 and identification of its response to azalomycin F by production of the red pigment carviolin. **A:** Map of Germany with marked location of the origin of soil sample and workflow for isolation of fungi. **B:** Carviolin reference, mono- and co-cultivation of *Penicillium* isolate 27 with *S. iranensis* WT and the azalomycin F-deficient mutant strain *S. iranensis* Δ*azlH* (left and right). Extracted ion chromatograms for carviolin (*m/z* 299 [M–H]^-^) derived from LC-MS analysis of culture extracts.

Sequencing of the ITS regions 1 and 2 suggested that all tested fungal soil isolates belong to the genus *Penicillium* (Supplementary Figure 10). This finding expands the group of signal-decoding fungi by the genus *Penicillium*, which is well known for its capability to produce NPs (*35*). Collectively, our data indicate the presence of widespread microorganism cooperatives wherein bacteria biosynthesize arginine-derived polyketides and fungi decode this molecular signal.

## Discussion

### Arginine-derived polyketides induce interkingdom microbial interactions

Given that healthy microbial communities play an enormous role in the health of the greater ecosystem (*3*), it is an important task to identify factors triggering interkingdom microbial interactions in microbial communities. Obvious candidates for such factors are NPs, produced by numerous microorganisms of prokaryotic and eukaryotic origin (*1, 10*). We have previously demonstrated the involvement of NPs in a specific bipartite interaction between the soil bacterium *Streptomyces rapamycinicus* and its closest relative *S. iranensis* with the fungus *Aspergillus nidulans* (*14*). This interaction leads to the activation of silent fungal NP BGCs including the production of orsellinic acid and derivatives thereof. What has been missing is the identification of widespread universal communication molecules that trigger bacterial-fungal cross-kingdom interactions in nature and help define the structure of microbial communities.

Based on the bipartite microbial interaction system between *S. iranensis* and *A. nidulans*, we here report the discovery of such a molecular bacterial signal *i.e.* arginine-derived polyketides, supported by genetic approaches, LC-MS-based metabolomics, transcriptome and proteome analyses, interaction studies and ecological investigations including the isolation of novel bacterial and fungal strains. These low molecular weight compounds produced by distantly related actinomycetes share the conserved ability to induce the production of NPs in phylogenetically diverse fungi (Figure 4B, Supplementary Figure 10, 11). Our results reveal that arginine-derived polyketides have a major impact on their surrounding microorganisms and thus shape their composition (Figure 4B). They directly impact surrounding microorganisms by inducing the production of fungal NPs that themselves impact other microorganisms. For example, *A. nidulans* responds with the production of orsellinic and lecanoric acid, which were shown to be produced by lichens (*36*). In line with a potential effect of marginolactones to promote symbioses is the finding of adverse effects of marginolactones on the green alga *Chlamydomonas reinhardtii.* At sublethal concentrations azalomycin F leads to the formation of a novel multicellular structure named gloeocapsoid that confers some protection to algal cells (*22*) and, most interestingly here, azalomycin F triggers green algae to accumulate and thereby hide in fungal mycelia from the adverse effects of azalomycin F (*16*) (Figure 4B). The compounds thus shape a lichen-like association between *C. reinhardtii* and *A. nidulans* that might have contributed to the evolution of lichens consisting of fungi and algae (*22*). Further, they also have the potential to impact the spatial partitioning of microrganisms in a microbial consortium. The induced fungal compound lecanoric acid has been found to specifically inhibit the growth of the plant-pathogenic basidiomycete *Rhizoctonia solani* (*37*). These compounds suggest a possible mechanism by which certain microorganisms are included in and excluded from a microbial consortium (Figure 4B).

**Fig. 4:**
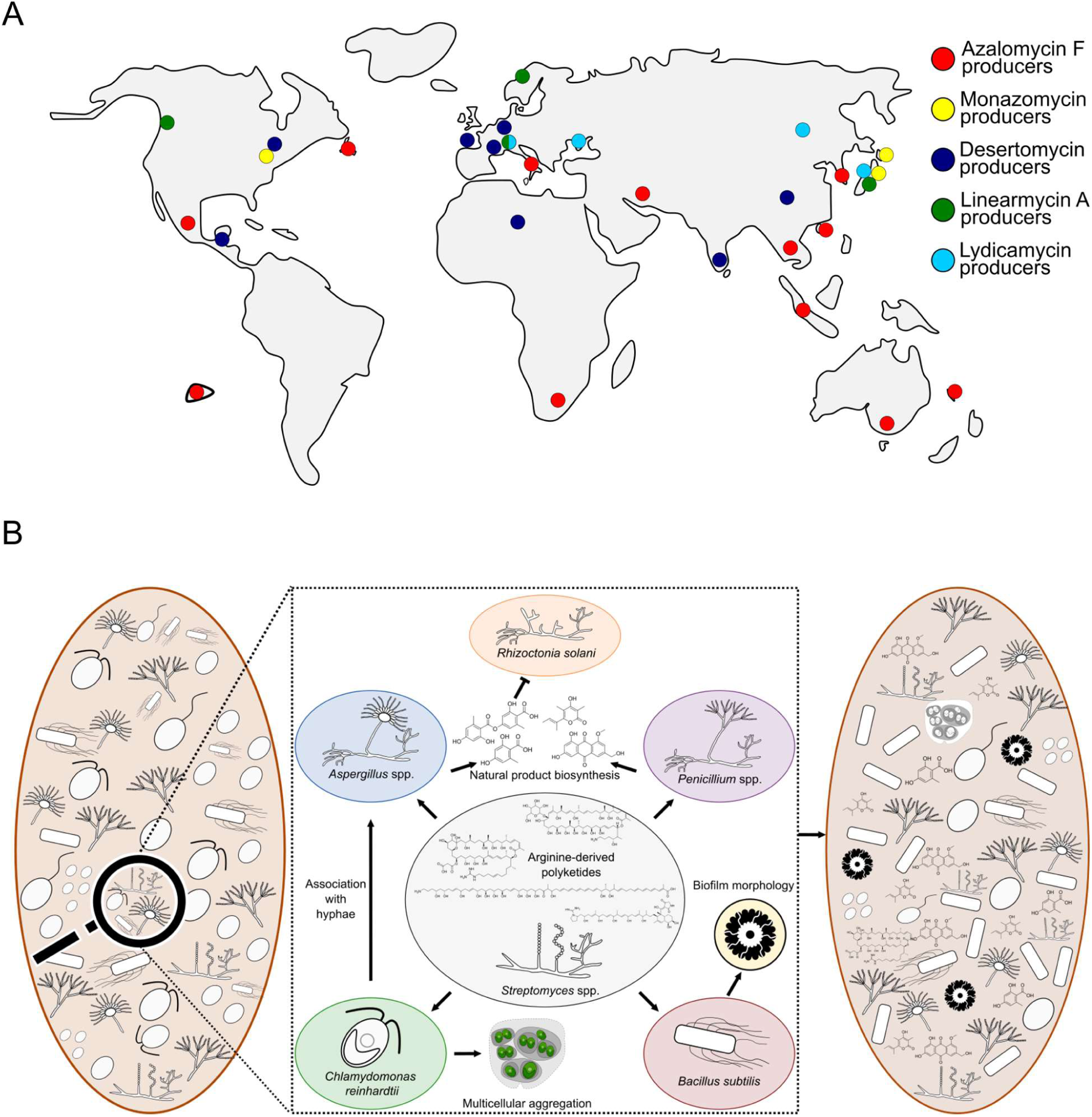
World-wide occurrence of arginine-derived polyketide producers and graphic summary of their effects on microorganisms and the structure and shape of a microbial consortium. **A:** World map without Antarctica indicating the place of isolation of bacteria producing the indicated arginine-derived polyketides based on genome and literature analyses. **B:** Graphic summary of the versatile roles played by arginine-derived polyketides and the produced fungal NPs in response on structure and shape of a microbial consortium.

Marginolactones can be divided into guanidyl-marginolactones such as azalomycin F, which are characterized by a guanidyl-moiety in their side chain, and amino-marginolactones such as desertomycin A and monazomycin, which contain a side chain terminal amino group formed from an arginine-derived guanidyl-moiety by a cluster-encoded agmatinase (*26, 38*). Similarly, linear arginine-derived polyketides also start with arginine as a biosynthetic precursor; however, they are not circularized (*24*). The amino- or guanidyl moiety seems to be essential for arginine-derived polyketides to act as signal molecules, since oasomycin B without such a group lost its inducing activity and had no antibacterial activity unless a positively charged moiety was reintroduced to the molecule (*29*).

As some marginolactones have been shown to bind to the membrane (*16, 39*), it was conceivable that their effect is connected to membrane damage. However, the membrane-damaging compounds amphotericin B or voriconazole, as well as compounds disturbing the fungal cell wall like caspofungin, did not have inducing activity, highlighting the specificity of marginolactones and that simple antifungal activity is not a trigger for the induction of the NP biosynthesis.

### Arginine-derived polyketides are ubiquitous signals for microorganisms

We herein show that arginine-derived polyketides serve as targeted signal molecules, produced by bacteria and decoded and further processed by fungi. Furthermore, we suggest that this is a universal signalling system, evidenced by the straightforward co-isolated of bacteria-fungal pairs and the fact that producing bacteria can be found on virtually all continents. The producing bacteria represent phylogenetically diverse streptomycetes (Supplementary Figure 11), suggesting that production of this signal molecule is advantageous for a number of different bacteria and that these molecules constitute a common signal of these bacteria. The presence of arginine-derived polyketides in soil is also supported by the observation that our sensor strain responded to soil supernatant. The receiver fungi thus far isolated belong to the genera *Penicillium* and *Aspergillus*, genera widespread in nature including in soil. These findings indicate a widespread phylogenetic as well as a global distribution of this type of communication. Furthermore, it is worth noting that not only fungi respond to arginine-derived polyketides, but that these compounds might also have signaling function on bacteria. For example, linearmycin A activates a two-component system of *Bacillus subtilis* resulting in induction of a specific biofilm morphology (*40, 41*).

Together, the ubiquitous distribution of actinomycetes producing these compounds on virtually all continents and the ease with which fungi decoding this chemical signal are isolated from soil suggests that arginine-derived polyketides represent a universally used component of the microbial communication network shaping microbial communities.

## Acknowledgments

We thank Dr. Amelia Barber for bioinformatics support. Christina Täumer is acknowledged for excellent technical support. This work was funded by the Deutsche Forschungsgemeinschaft (DFG, German Research Foundation) – Project-ID 239748522 – CRC 1127 ChemBioSys, the Cluster of Excellence Balance of the Microverse under Germany’s Excellence Strategy - EXC 2051 - Project-ID 390713860 and the DFG Collaborative Research Center/Transregio FungiNet 124 ‘Pathogenic fungi and their human host: Networks of Interaction’ (project Z2; project number 210879364).

## Author contributions

Conceptualization: AAB, CH, MKCK, MCS, VS

Methodology: JB, MR, MCS, MKCK, AJK, CH, TN, OK, TK, VS

Investigation: JB, MKCK, TN, MCS, MR, LZ, TK, AJK, VS

Visualization: LZ, MCS, MKCK, VS, JB

Funding acquisition: AAB, CH

Project administration: AAB

Supervision: AAB, CH

Writing – original draft: AAB, MKCK

Writing – review & editing: AAB, CH, MKCK, MCS, VS

## Competing interests

Authors declare that they have no competing interests.

## Data availability statement

Proteome, transcrptome, genome, and ITS sequences will be made accessible for the final publication.

## Materials and Methods

### Microorganisms, plasmids, media and cultivation

All microbial strains and plasmids used in this study are listed in Supplementary Table 4. Primers used are listed in Supplementary Table 5.

### Cultivation of microorganisms

*Streptomyces iranensis* DSM41954 (HM35^T^) wild type and deletion mutants, *Streptomyces macronensis* UC 8271 (NRRL12566) and *Streptomyces mashuensis* DSM40896 were grown as described in (*43*). *Streptomyces* spp. soil isolates 7, 45, 48, 102, 124, 176, 219, and 280 were inoculated with 2.5 × 10^7^ spores in TSBY in Erlenmeyer flasks with cotton wool plugs and incubated at 28 °C and 180 rpm for 3 days. Spores were generated by streaking 200 – 300 µL of densely grown cultures on oatmeal agar plates, which were incubated at 28 °C for 14 days and spores harvested.

*Aspergillus nidulans* RMS011 and *Aspergillus fumigatus* ATCC 46645 were cultivated as described in (*44*).

### Co-cultivation of *Aspergillus* spp. with *Streptomyces* spp

4-days-old pre-cultures of *Streptomyces* spp. were set-up as described above. Mycelia of *A. fumigatus* or *A. nidulans* overnight cultures (∼16 h old) in AMM were separated from the medium using Miracloth (Merck Millipore, Darmstadt, Germany) and placed in fresh AMM (in the case of *A. nidulans* supplemented with 5 mM L-arginine, 1 mL/L trace elements, 0.3 mM FeSO4 and 3 µg/mL PABA) (*45*). Either 1/20 volume of the streptomycete culture (*46*) or 5 - 40 µg/mL purified arginine-derived polyketide (desertomycin A, monazomycin A, azalomycin F, linearmycin A, lydicamycin & oasomycin B) were added to the culture, which was then further incubated at 37 °C with shaking at 200 rpm. Desertomycin A, lydicamycin, and monazomycin were purchased from Santa Cruz Biotechnology (Dallas, USA), linearmycin A & oasomycin B were purchased from BioAustralis (Sydney, Australia). Azalomycin F was purified from *S. iranensis* as described below. Samples for LC-MS analyses were taken after 12 h (for *A. fumigatus*) and 24 h (for *A. nidulans*) of co-cultivation and extracted as described below.

### Isolation of filamentous bacteria from soil

Soil at the surface was collected near the castle ruin “Eutinger Tal” in Eutingen im Gäu, Baden-Württemberg, Germany (GPS: 48.4661535, 8.7291989) on June 24^th^, 2021. This site was chosen due to a lack of agricultural and forestry as a consequence of its status as protected landscape. To the best of our knowledge, this site has never been subject to isolation of microorganisms before and hence no sampling bias was predicted. In total, approximately 600 mg of soil were used to isolate filamentous bacteria. On three separate occasions, roughly 200 mg of soil were mixed with 3 mL PBS and vortexed intensively. The samples were allowed to sediment for 1.5 – 2 h and the supernatants were transferred to new tubes and heated for 10 min at 50 °C to induce germination of spores. The samples were allowed to cool down for 10 minutes at room temperature. Undiluted samples and samples diluted 1:100 in PBS were streaked on TAP-Agar + 100 µg/mL cycloheximide and 50 µg/mL nalidixic acid to inhibit growth of fungi and Gram-negative bacteria. The agar plates were subsequently incubated at 28 °C for 7 – 14 days. Single colonies that showed filamentous growth were re-streaked on oatmeal agar. After these microorganisms had re-grown, colonies were inoculated into 3 mL GYM, 3 mL TSB, and 3 mL M79 medium (*47*). As soon as the cultures had grown to a high density, they were cocultured with *A. nidulans* strain *orsA*p-nLuc-GFPs to evaluate their ability to activate the *ors*-BGC. GFPs- inducing bacteria were cocultured with *A. nidulans* RMS011 and production of orsellinic acid and derivatives was evaluated by HPLC-MS. Bacteria inducing fungal orsellinic acid production were subject to genome sequencing using Illumina NextSeq 2000 (Paired-end sequencing, read length 150 nt, 100 × coverage, 10 M reads, StarSeq GmbH, Mainz, Germany). Genomic DNA was isolated using the NucleoSpin Microbial DNA Mini kit (Macherey & Nagel, Düren, Germany) according to the manufacturer’s manual. Potential natural product biosynthesis gene clusters were identified using antiSMASH (*48*) and the BLAST algorithm (*49*). Whole genome-based phylogenetic analysis was carried out using the Type Strain Genome Server (TYGS) provided by the DSMZ, Germany (*50*). Data are will be made accessible for the final publication.

### Isolation of filamentous fungi from soil

For the isolation of filamentous fungi, the same soil sample was used as for the isolation of bacteria. 200 mg of soil were mixed with 3 mL PBS and vortexed intensely. The sample was allowed to sediment for 1.5 – 2 h. Undiluted and samples diluted 1:10 and 1:100 were plated on malt extract agar plates (20 g/L malt extract, 2 g/L yeast extract, 10 g/L glucose, 0.25 g/L NH4Cl, 0.25 g K2HPO4, 20 g Agar, pH = 6) supplemented with 50 µg/mL nalidixic acid and 25 µg/mL kanamycin to inhibit growth of bacteria. The agar plates were incubated at 28 °C for 3 – 17 days. Single colonies of fungi were re-streaked on AMM agar containing Hutner’s trace elements (*51, 52*). Spores were harvested with 5 mL 0.9 % (w/v) NaCl and stored at - 80 °C in 50 % (v/v) glycerol. For screening for their response to azalomycin F, the fungal isolates were co-cultivated with *S. iranensis* or 10 µg/mL azalomycin F in 24-well plates in a Thermo-shaker PST-60HL (Biosan, Riga, Latvia). For this purpose, 1 mL AMM-Hutner’s medium was inoculated with 10^6^ - 10^7^ spores and they were cultivated for 2 days at 28 °C and 600 rpm. When fungal growth was observed, the medium was replaced by fresh AMM medium supplemented with Hutner’s trace elements and 50 µL of an *S. iranensis* culture or 10 µg/mL azalomycin F were added to the fungal mycelium. The coculture was incubated for 1 – 2 days at 28 °C and 600 rpm. All isolates showing a visible color-change when cocultured with *S. iranensis* or azalomycin F compared to the fungus alone were further investigated. Candidate fungal isolates were pre-cultured in 50 mL AMM supplemented with Hutner’s trace elements and cocultured in 15 mL fresh AMM supplemented with Hutner’s trace elements with 750 µL of cultures of *S. iranensis* WT or *S. iranensis* Δ*azlH*. Formation of natural products was evaluated by HPLC-MS after 1 – 2 days. To determine the genus of isolated fungi, their gDNA was isolated using the method described in Schroeckh *et al*. (*46*). Then, their ITS1 and ITS2 regions flanking the 5.8S rDNA were amplified by PCR using proof-reading Phusion™ High-Fidelity DNA Polymerase (Thermo Fisher Scientific, Dreieich, Germany) and the primers ITS1 and ITS4 (*53*). Sequencing was carried out by LGC Genomics (Berlin, Germany) and the obtained sequences were analyzed by using the BLAST algorithm and MEGA X (*49, 54*). The carviolin standard used for comparison with extracts of *Penicillium* sp. 27 was purchased from Merck, Darmstadt, Germany. ITS sequences will be made accessible for the final publication.

### Extraction and detection of natural products

Extraction and detection of natural products were carried out as described in Stroe et al. (*44*). For measurement of azalomycin F in the biomass and in the supernatant of an *A. nidulans* culture, the biomass was separated from the medium using Miracloth (Merck Millipore, Darmstadt, Germany). Identification of desertomycin A, monazomycin, lydicamycin, and linearmycin A was achieved by comparison with authentic references purchased from Santa Cruz Biotechnology (Dallas, USA) and BioAustralis (Sydney, Australia).

Matrix-assisted laser desorption/ ionization mass spectrometry (MALDI Imaging MS) was performed as previously described in Krespach *et al* (*43*).

### Production and purification of azalomycin F from *S. iranensis*

The azalomycin F complex from *S. iranensis* was produced and purified as previously described in Krespach et al. (*43*).

### Generation of *S. iranensis* deletion strains

Gene deletions in *S. iranensis* were generated as described (*55*) and verified by Southern blot analysis (Supp. Fig. 13), essentially according to Southern (*56*). Bacterial genomic DNA was isolated using the NucleoSpin Microbial DNA Mini kit (Macherey & Nagel, Düren, Germany). Oligonucleotide sequences used are listed in Supplementary Table 5. For Southern blot analysis enzymes used to cleave chromosomal DNA were *Bam*HI (New England Biolabs, Frankfurt, Germany) for verification of *ΔbldA*, *Bst*EII (New England Biolabs, Frankfurt, Germany) for Δ*bldD*, and *Pst*I (New England Biolabs, Frankfurt, Germany) for *ΔbldH*.

### Generation of the *A. nidulans orsA*p-nLuc-GFPs reporter strain and screening for bacterial isolates inducing GFPs-dependent fluorescence

To study the activation of the *ors* BGC in *A. nidulans*, the human codon-optimized nanoluciferase (nLuc) gene (Promega, Mannheim, Germany) and the GFPspark (GFPs) were translationally fused and placed under the control of the native *orsA* promoter by integrating the gene fusion and the *pabaA* gene as selection marker *in locus* upstream of the *orsA* gene. The transformation cassette was constructed as previously described (*57*). Approximately 2000 bp sequences homologous to the regions upstream and downstream of *orsA* (*AN7909*) were amplified and assembled with GFPs, nLuc and the *pabaA* gene into pUC18 using the NEBuilder HiFi DNA Assembly Master Mix (New England Biolabs, Frankfurt, Germany) according to the manufacturer’s instructions. Transformation of *A. nidulans* with the PCR-amplified plasmid was carried out as described before (*58*). Colonies of transformant strains were selected on AMM agar plates supplemented with 5 mM L-arginine, 1 mL/L trace elements, and 0.3 mM FeSO4. The genomic structure of the transformant strain was verified by Southern blot analysis as described in (*56*) using a probe directed against the *orsA* gene (Supplemental Figure 6), which was amplified using primers MM031 and MM065 (Supplemental Table 5).

For evaluation of the ability of soil-isolated bacteria to activate *orsA* indicated by green fluosrescence and activity of the nanoluciferase, the *A. nidulans orsA*p-nLuc-GFPs strain was cultured in AMM supplemented with 5 mM L-arginine, 1 mL/L trace elements, and 0.3 mM FeSO4. Mycelia of the overnight culture (∼16 h old) in AMM were separated from the medium using Miracloth (Merck Millipore, Darmstadt, Germany) and distributed in a 24-well plate (Greiner bio-one, Cellstar, Merck, Darmstadt, Germany) with 1 mL fresh AMM supplemented with 5 mM L-arginine, 1 mL/L trace elements, and 0.3 mM FeSO4. After 6 h of incubation at 37 °C and shaking at 600 rpm in a tabletop shaker (Thermo-shaker PST-60HL, Biosan, Riga, Latvia), fluorescence and brightfield images were taken on a Keyence BZ-X800 microscope (Keyence, Osaka, Japan) with 4× magnification.

For quantification of the nLuc activity, the fungal mycelium was first added to a 2 mL screwcap tube, filled with Zirconia beads (Thermo Fisher Scientific, Waltham, USA) and homogenized by 2 rounds of 30 s in a Fast Prep (Tabletop Speedmill Plus, Analytic Jena, Jena, Germany). After centrifugation, the luminescence of the supernatant was analyzed using the Nano-Glo luciferase assay system (Promega, Mannheim, Germany) according to the manufacturer’s instructions, using a microtiter plate reader (TECAN, Männedorf, Switzerland).

### RNA isolation, cDNA library construction and sequencing

Total RNA of *S. iranensis* wild type and its deletion mutants was isolated using the Direct-zol™ RNA MiniPrep Plus Purification Kit (Zymo Research Europe, Freiburg, Germany). Samples were taken after 48 h of cultivation in TSB medium. Bacterial cells were disrupted by bead beating for 4 min in a Fast Prep (Tabletop Speedmill Plus, Analytic Jena, Jena, Germany). DNAse treatment using Baseline-ZERO DNAse (Lucigen, Middleton, WI, USA) was followed by an RNA Clean & Concentrator-5 (Zymo Research Europe, Freiburg, Germany) clean-up procedure. In each case, total RNA from three replicates was pooled and 2-3 µg of RNA were processed for the library preparation. Library construction, Illumina NextSeq 500 paired end sequencing, mapping and normalizing of the reads were performed by StarSEQ GmbH (Mainz, Germany). Reads were aligned to the NCBI reference genome for *S. iranensis* (Assembly GCA_000938975.1). Transcripts were normalized by counting the number of transcripts per million (TPM) (Wagner, Kin et al. 2012). The RNA-Seq data will be made accessible for the final publication.

### Protein purification from *S. iranensis* and proteomics

Protein isolation and LC-MS/MS analysis for identification of proteins were performed essentially as previously described in Stroe et al. (*44*), Mass spectrometry analysis was performed on a Q Exactive Plus instrument (Thermo Fisher Scientific) at a resolution of 140,000 FWHM for MS1 scans and 17,500 FWHM for MS2 scans. Tandem mass spectra were searched against the NCBI database of *Streptomyces iranensis* (2019/01/24). A strict false discovery rate (FDR) < 1% (peptide and protein level) and at least a search engine threshold >30 (Mascot), >4 (Sequest HT) or >300 (MS Amanda) were required for positive protein hits. Label-free protein quantification was based on the Minora algorithm of PD2.2 using a signal-to-noise ratio >5. The mass spectrometry proteomics data will be made accessible for the final publication.

### qRT-PCR

Expression of the *orsA* gene was quantified as previously described (*46*) using primers orsA_FW and orsA_RV. qRT-PCR results were analyzed by means of the QuantStudio^TM^ Design & Analysis software (ver. 1.5.2; Applied Biosystems, Foster City, CA). Relative gene expression was calculated with the ΔΔCt method, normalized to the expression of the *A. nidulans* γ-actin gene *AN6542* as internal standard (primers actin FW and RV) using the formula 2^−(Ct *orsA* − Ct AN6542)^ and compared to an *A. nidulans* monoculture as calibrator.

## Supplementary Tables and Legends to Supplementary Figures

**Supplementary Table 1:** Comparison of the lydicamycin BGC of *Streptoymces* sp. ID38640 with sequences of bacterial soil isolate 219. Homologous genes were found in bacterial soil isolate 219 for each gene of the lydicamycin BGC in *Streptomyces* sp. ID38640 (*59*).

| Lydicamycin BGC access numbers | Percentage of identical nucleotides of bacterial soil isolate 219 to *Streptomyces* sp. ID38640 (59). | Location |
| --- | --- | --- |
| G7Z12_32710 (4-Guanidinobutyryl-CoA biosynthesis) | 97% | Contig 34 |
| AVV61973.1 | 97% | Contig 34 |
| AVV61974.1 | 97% | Contig 34 |
| AVV61975.1 | 97% | Contig 34 |
| AVV61976.1 | 98% | Contig 34 |
| AVV61977.1 (4-Guanidinobutyryl-CoA loading) | 97% | Contig 34 |
| AVV61978.1 | 98% | Contig 34 |
| AVV61979.1 | 96% | Contig 34 |
| AVV61980.1 | 94% | Contig 28<br>Contig 240<br>Contig 241<br>Contig 242 |
| AVV61981.1 | 95% | Contig 28 |
| AVV61982.1 | 96% | Contig 28 |
| AVV61983.1 | 95% | Contig 28 |
| AVV61984.1 | 97% | Contig 28 |
| AVV61985.1 | 97% | Contig 28 |
| AVV61986.1 | 98% | Contig 28 |
| AVV61987.1 | 98% | Contig 28 |
| AVV61988.1 | 98% | Contig 28 |
| AVV61989.1 | 97% | Contig 28 |
| AVV61990.1 | 97% | Contig 28 |
| AVV61992.1 | 98% | Contig 28 |
| AVV61993.1 | 97% | Contig 28 |
| AVV61994.1 | 97% | Contig 5 |
| AVV61995.1 | 97% | Contig 5 |
| AVV61996.1 (4-Guanidinobutyric acid biosynthesis) | 97% | Contig 5 |
| AVV61997.1 | 98% | Contig 5 |

**Supplementary Table 2:**
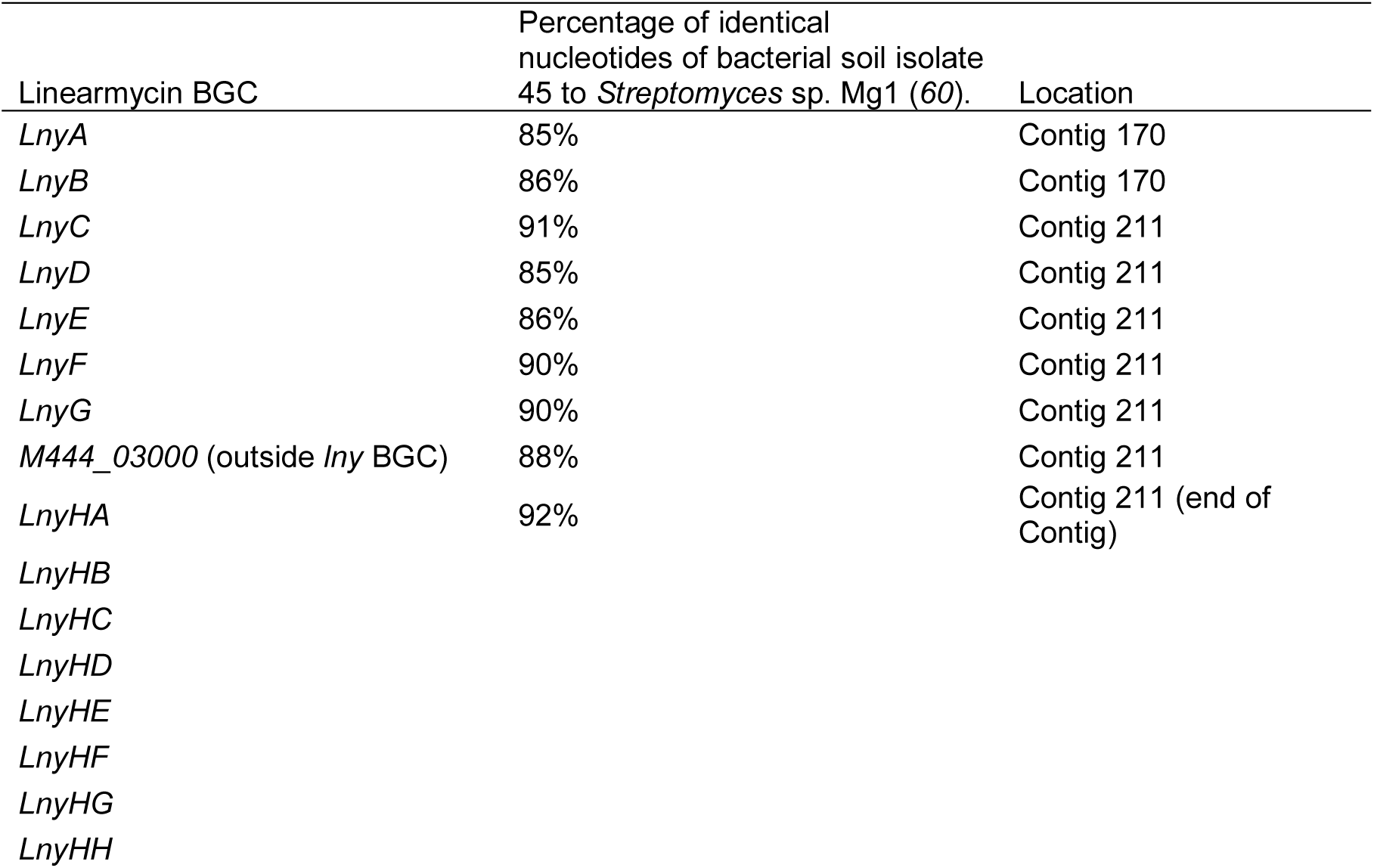
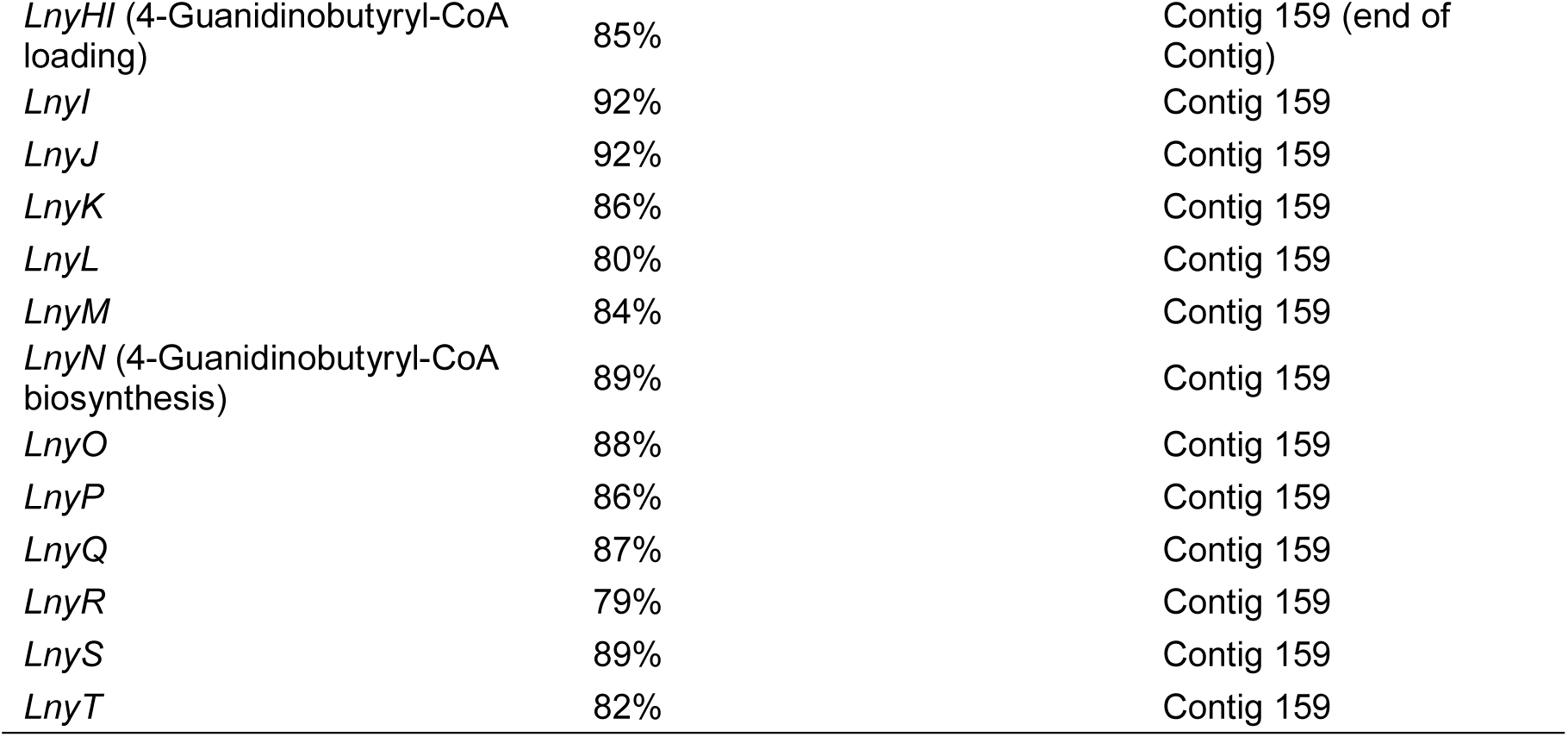
Comparison of the linearmycin BGC of *Streptomyces sp*. Mg1 (*60*) with sequences of bacterial soil isolate 45. All genes encoding the tailoring enzymes of the biosynthesis of linearmycin were found to be highly conserved in bacterial soil isolate 45 (*lnyA*-*G* and *lnyI*-*T*). Homologs of the polyketide synthase genes *lnyHA* and *lnyHI* were also identified. The core polyketide synthase genes could not be assigned to specific contigs since their repetitive sequences were not resolved by illumina sequencing.

**Supplementary Table 3:** World-wide occurrence of arginine-derived polyketide-producing actinomycetes.

| Compound | Producing microorganisms | Location of isolation | Source |
| --- | --- | --- | --- |
| Desertomycin A | *Streptomyces flavofungini* | Desert sand, Africa | (61) |
|  | *Streptomyces spectabilis* BT352 | Southern France | (62) |
|  | *Streptomyces macronensis* | Belize | (63, 64) |
|  | *Streptomyces nobilis* JCM4274 (*Streptomyces* sp. 725) | Hesbaye, Belgium | (65, 66) |
|  | *Streptomyces alboflavus* SC11 | Western Sichuan Plateau, China | (67) |
|  | *Streptomyces baldacii* subsp. *netropsis* (*Streptomyces* sp. DSM5990) | India | (68, 69) |
| Desertomycin G | *Streptomyces althioticus* MSM3 | Surface of seaweed *Ulva* sp., Cantabrian Sea | (70) |
| Monazomycin/<br>Takacidin | *Streptoverticillium griseoverticillatum* | Iwate Prefecture, Japan | (71) |
|  | *Streptomyces mashuensis* | Lake Mashu, Japan | (72) |
|  | *Streptoverticillium* sp. UC8633 | Indiana, USA | (73) |
| Azalomycin F | *Streptomyces iranensis* | Teheran, Iran | (74) |
|  | *Streptomyces rapamycinicus* | Rapa Nui | (75) |
|  | *Streptomyces malaysiensis*<br>MJM1968 | Imsil, South Korea | (76) |
|  | *Streptomyces* sp. 211726 | Mangrove tree<br>rhizosphere,<br>Wenchang, China | (77) |
|  | *Streptomyces violaceusniger*<br>Tü4113 | Otway National Park,<br>Australia | (78) |
|  | *Streptomyces* sp. M56 | Termite mount, South<br>Africa | (79) |
|  | *Streptomyces*<br>*melanosporofaciens* DSM40318 | Wheat Field near<br>Pescara, Italy | (80) |
|  | *Streptomyces rhizosphaericus*<br>NRRL B-24304 | Roots of<br>*Paraserianthes*<br>*falcataria*, Java,<br>Indonesia | (81) |
|  | *Streptomyces javensis*<br>DSM41764 | Roots of<br>*Paraserianthes*<br>*falcataria*, Java,<br>Indonesia | (81) |
|  | *Streptomyces sabulosicollis* | Sand dune, Indonesia | (82) |
|  | *Streptomyces endocoffeicus* | Coffea arabica roots,<br>Thailand | (83) |
|  | *Streptomyces aureoverticillatus*<br>HN6 | Hainan, China | (84) |
|  | *Streptomyces buecherae*<br>NA00687 | Hainan, China | (85) |
|  | *Streptomyces buecherae*<br>AC541<sup>T</sup> | Bat cave in Southern<br>Mexico | (85) |
|  | *Streptomyces solisilvae*<br>HNM0141 | Hainan, China | (86) |
|  | *Streptomyces* sp. 11-1-2 | Potato Scab,<br>Newfoundland,<br>Canada | (87) |
|  | *Streptomyces yatensis*<br>DSM41771 | New caledonia | (88) |
| Linearmycin A | *Streptomyces* sp. Mg1 | Mendenhall Glacier,<br>Alaska, USA | (89) |
|  | *Streptomycesnojiriensis*<br>JCM3382 | Nagano Prefecture,<br>Nojiri, Japan | (90, 91) |
|  | *Streptomyces* sp. ADI95-16 | Marine sponge *Antho*<br>*dichotoma*, Trondheim<br>Fjord, Norway | (92) |
|  | *Streptomyces* sp. 45 | Topsoil, Eutingen im Gäu, Germany | This study |
| Lydicamycin | *Streptomyces lydicus* 2249-S3 | Ohata-machi, Shimokita-gun, Aomori Prefecture, Japan. | (93) |
|  | *Streptomyces lydicamycinicus* TP-A0598<sup>T</sup> (=NBRC 110027<sup>T</sup>) | Deep sea water, Namerikawa, Toyama, Japan | (94, 95) |
|  | *Streptomyces* sp. strain Je 1–6 | Mount Koshka, Ukraine | (96) |
|  | *Streptomyces* sp. NEAU-S7GS2 | Saline-alkaline field in Durbert, Daqing, China | (97) |
|  | *Streptomyces platensis* TP-A0598 | Toyama Bay, Japan | (98) |
|  | *Streptomyces* sp. 7, 48, 102, 124, 176, 219, 280 | Topsoil, Eutingen im Gäu, Germany | This study |

**Supplementary Table 4:**
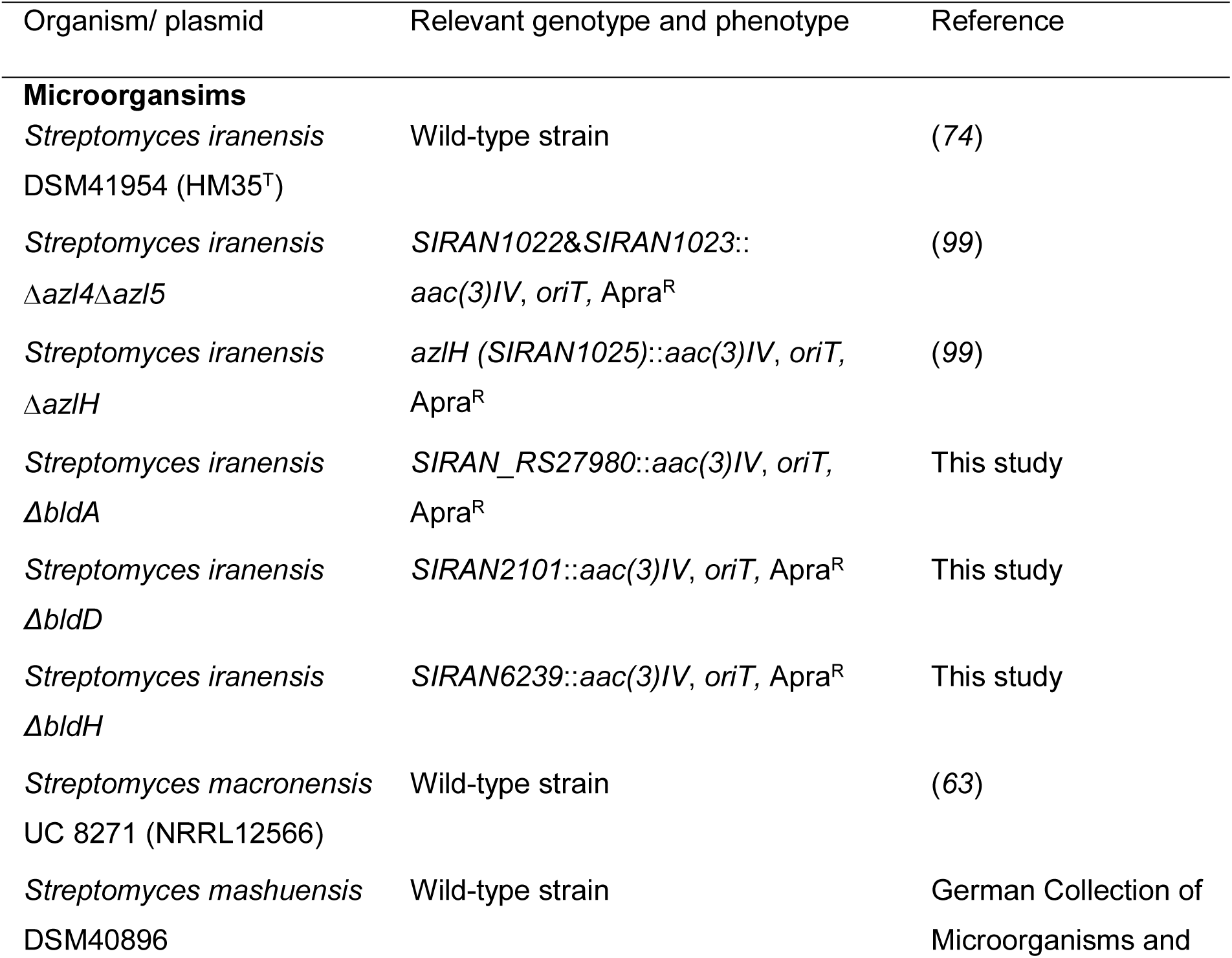
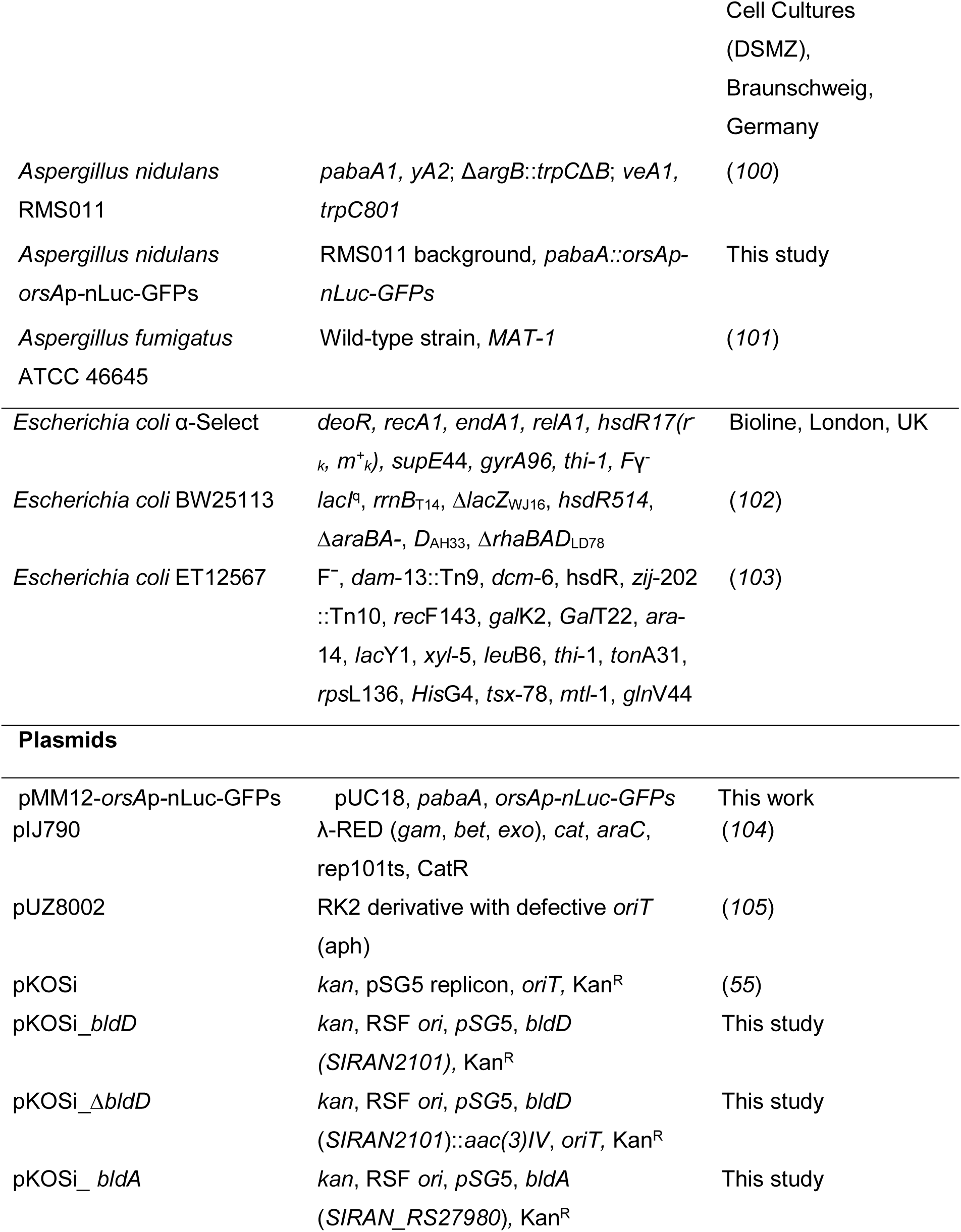
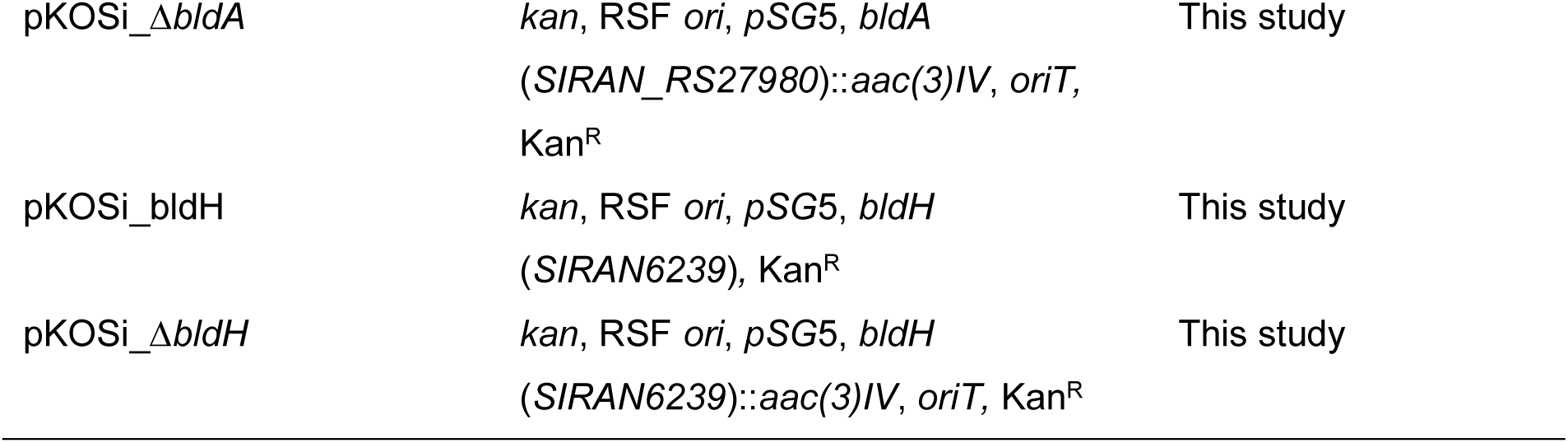
Microorganisms and plasmids, their genotype and reference.

**Supplementary Table 5:**
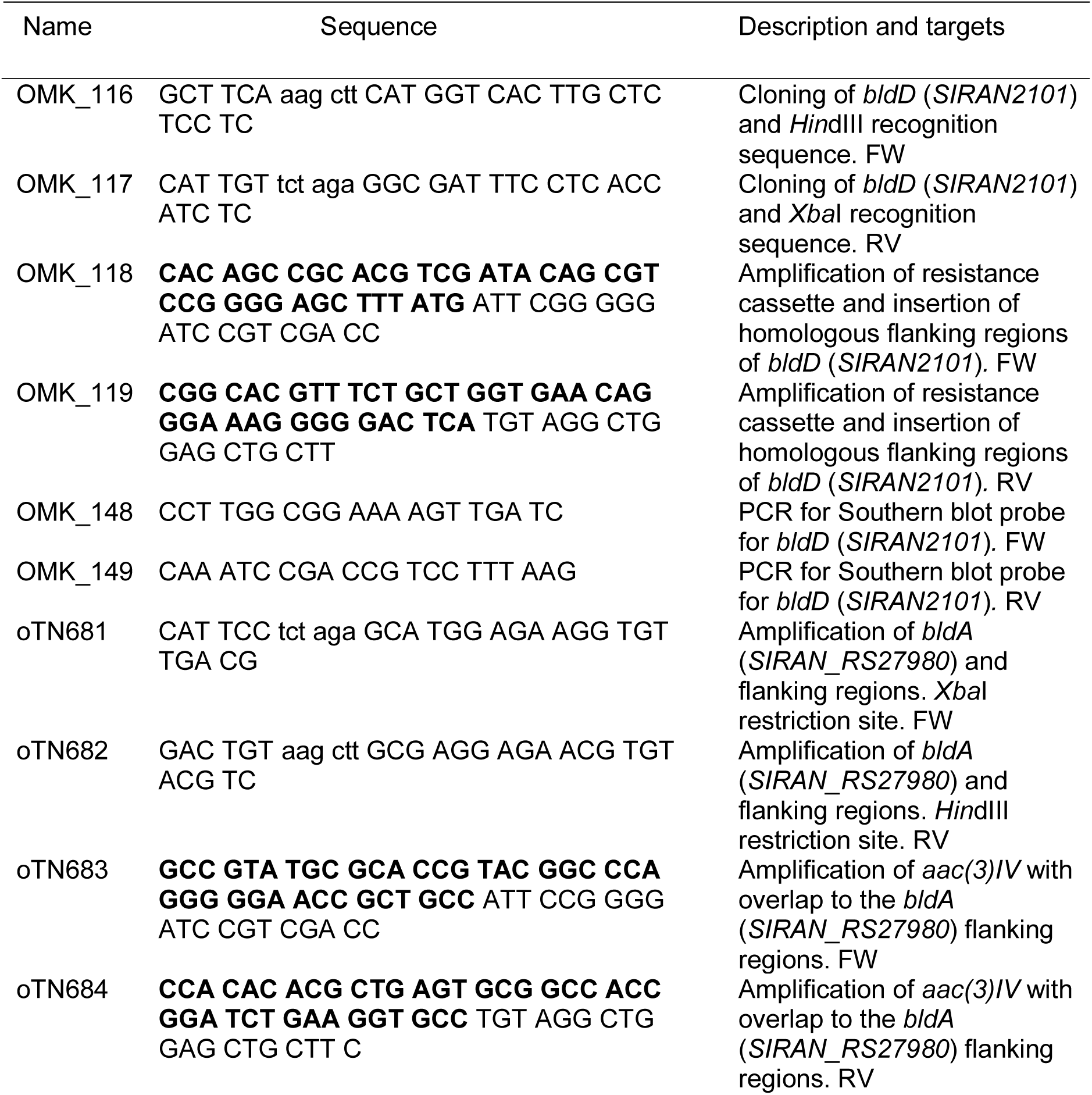
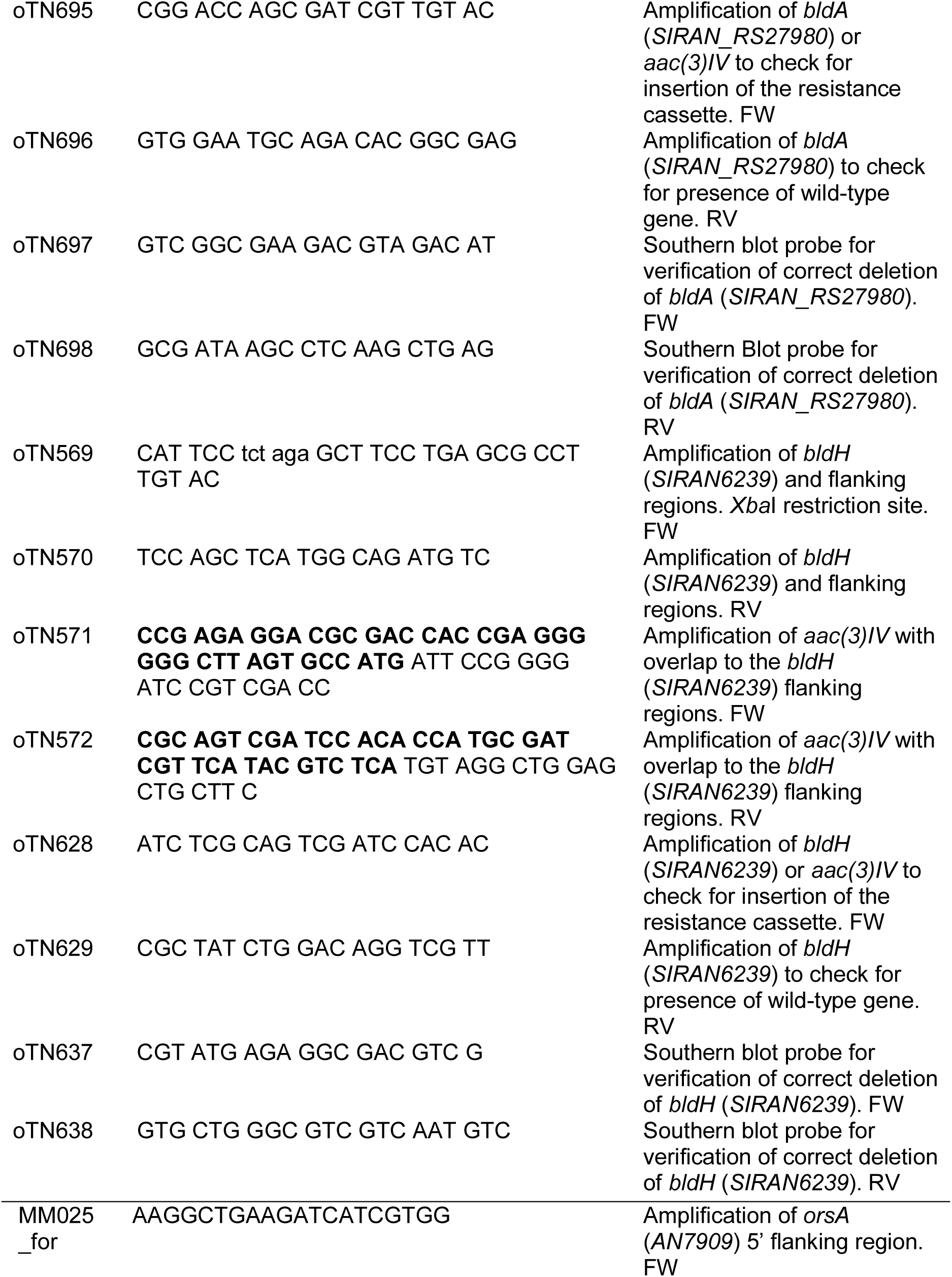
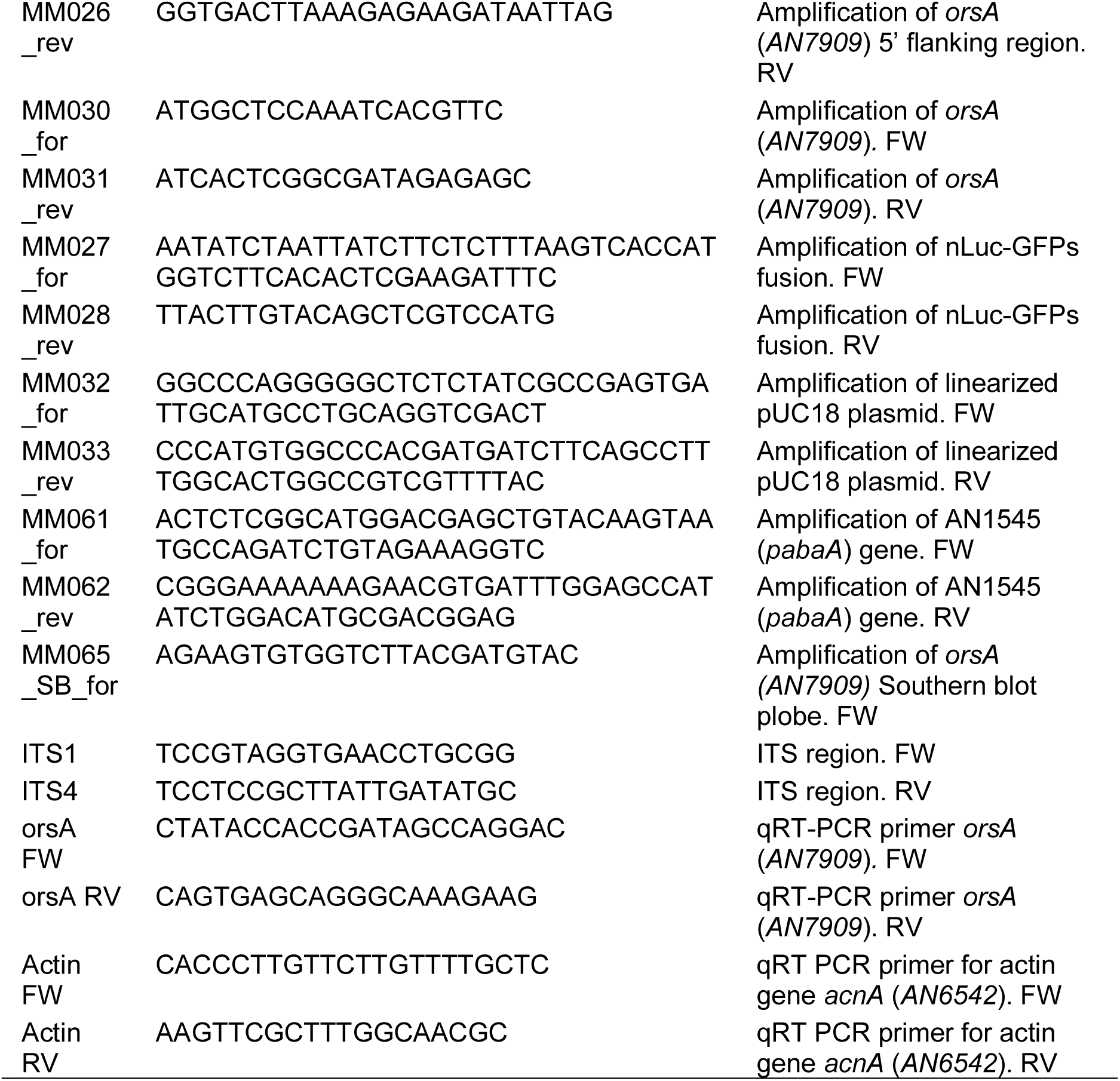
Primers used in this study.

## Supplementary Figures

**Supp. Fig.1:**
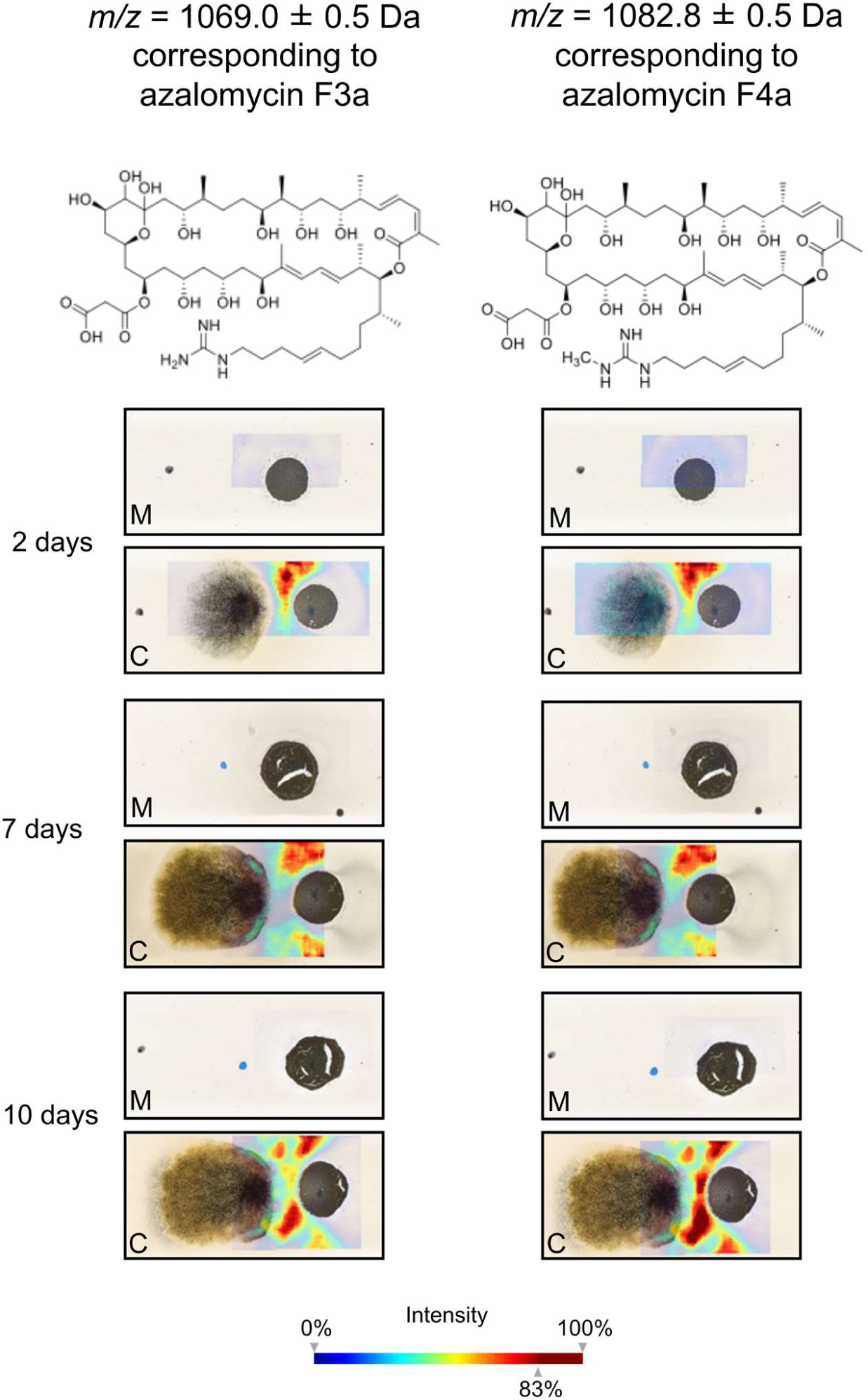
MALDI-IMS time course of cocultivation of *A. nidulans* with *S. iranensis*. Visualization of masses corresponding to the main azalomycin F derivatives azalomycin F3a (*/mz* 1069.0 ± 0.5) and azalomycin F4a (*m/z* 1082.8 ± 0.5) produced by *S. iranensis* (right on the glass slide) after 2, 7 and 10 days of cocultivation with *A. nidulans* (left on the glass slide). Abundances of the analyzed masses are depicted as a heat map from low abundance (blue) to high abundance (red). M, monoculture *S. iranensis*; C, coculture of *S. iranensis* with *A. nidulans*.

**Supp. Fig.2:**
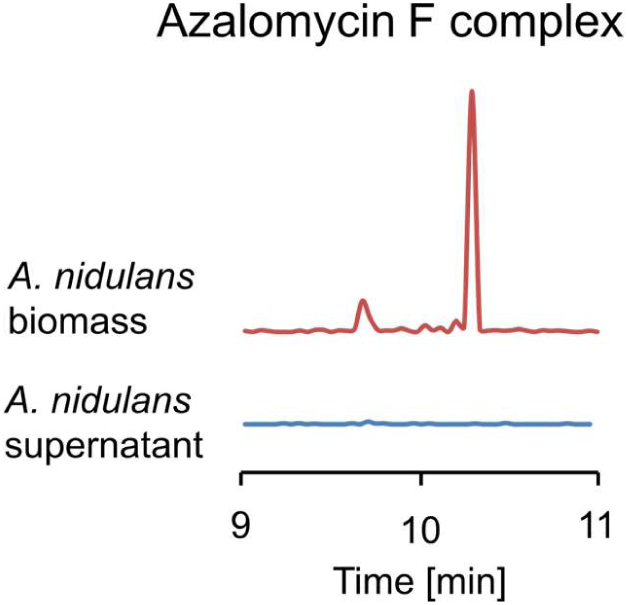
Extracted ion chromatogram of azalomycin F3a (*/mz* 1066 [M-H]^-^) derived from LC-MS analyses of biomass and culture supernatant of an *A. nidulans* culture supplemented with azalomycin F.

**Supp. Fig.3:**
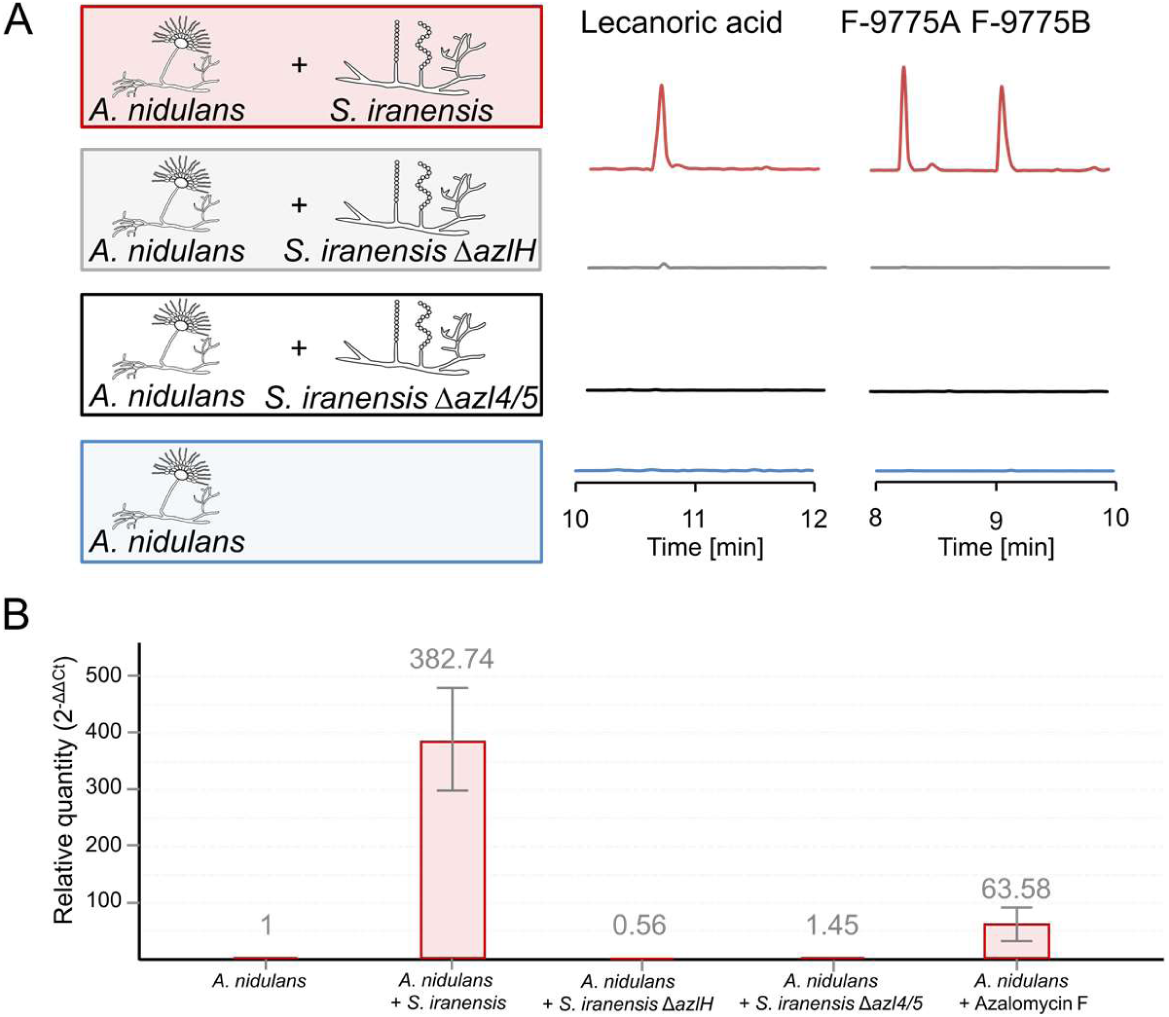
Analysis of orsellinic acid derivatives and expression of orsellinic acid biosynthesis gene *orsA* of *A. nidulans* during cocultivation of the fungus with *S. iranensis* or addition of azalomycin F to the culture medium. **A:** Coculture of *A. nidulans* with *S. iranensis* WT and mutant strains Δ*azlH and azl5/5* and extracted ion chromatogram of lecanoric acid *(m/z* 317 [M-H]^-^) and isobaric compounds F-9775A and F-9775B (*/mz* 395 [M-H]^-^) derived from LC-MS analysis of culture supernatant. **B:** qRT-PCR analysis of the orsellinic acid biosynthesis gene *orsA* of *A. nidulans* cocultivated with indicated *S. iranensis* strains or in monoculture supplemented with azalomycin F.

**Supp. Fig.4:**
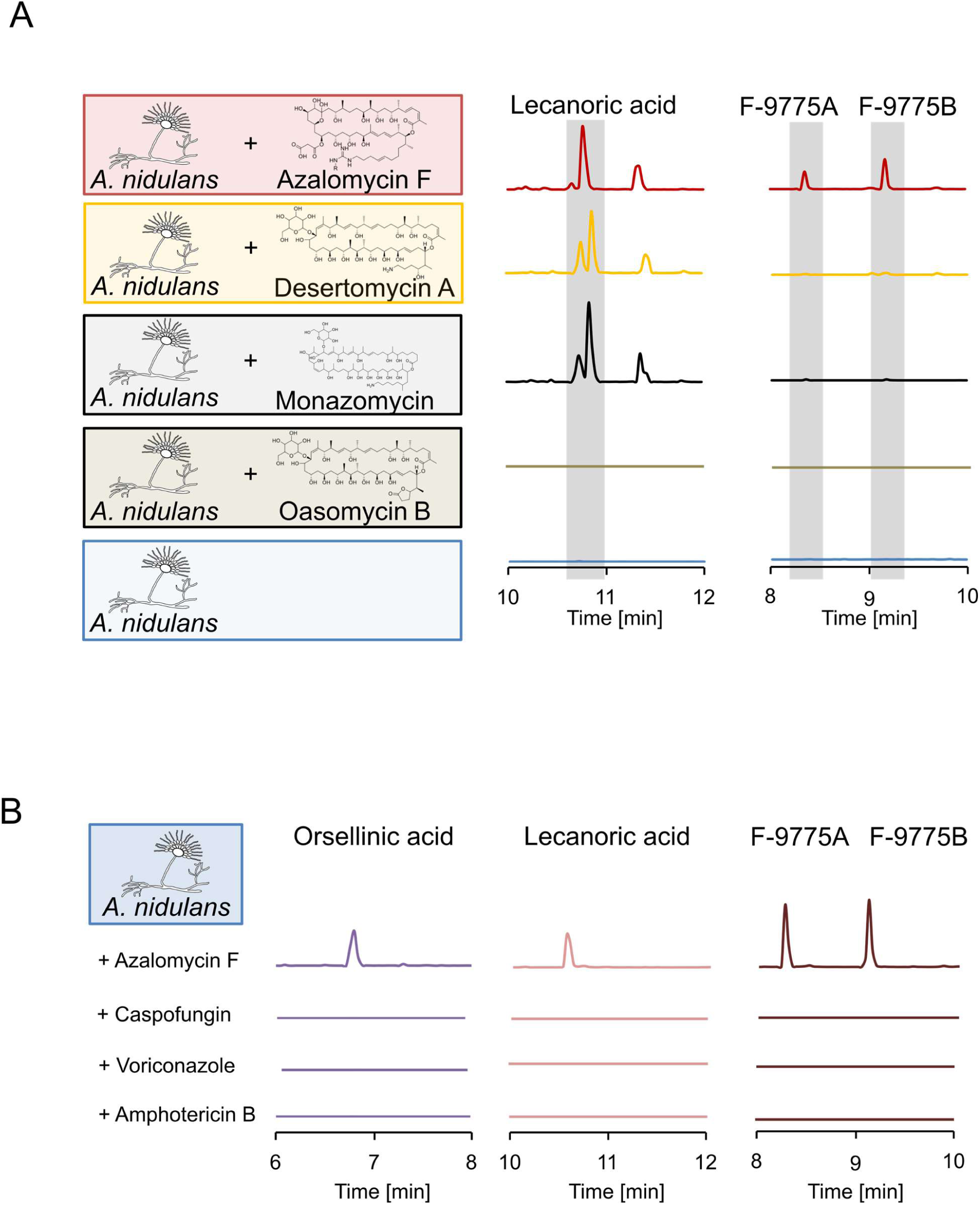
Analysis of margionolactones, derivatives thereof and membrane- and cell-wall disturbing compounds for their *ors* BGC inducing activity. **A:** Cultivation of *A. nidulans* supplemented with indicated compounds and extracted ion chromatograms for lecanoric acid (*/mz* 317 [M–H]^-^) and F-9775A and F-9775B (*m/z* 395 [M–H]^-^) derived from LC-MS analysis of culture supernatant. **B:** Cultivation of *A. nidulans* supplemented with indicated compounds and extracted ion chromatograms for orsellinic acid (*m/z* 167 [M–H]^-^), lecanoric acid (*/mz* 317 [M–H]^-^) and F-9775A and F-9775B (*m/z* 395 [M–H]^-^) derived from LC-MS analysis of culture supernatant.

**Supp. Fig.5:**
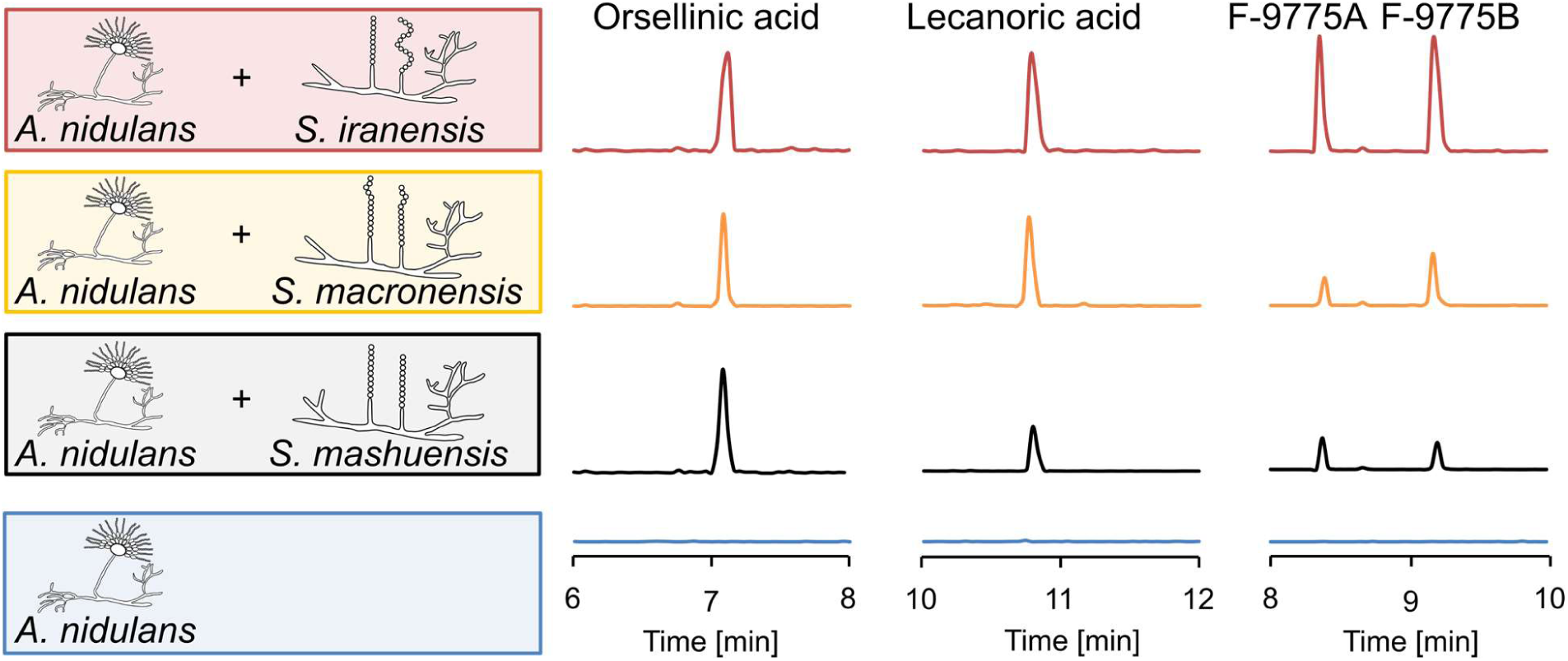
Production of *ors* compounds by *A. nidulans* triggered by marginolactone- producing streptomycetes. Cocultures of *A. nidulans* with indicated *Streptoymces* species and extracted ion chromatograms for orsellinic acid (*m/z* 167 [M–H]^-^), lecanoric acid (*/mz* 317 [M–H]^-^) and F-9775A and F-9775B (*/mz* 395 [M–H]^-^) derived from LC-MS analysis of culture supernatant.

**Supp. Fig.6:**
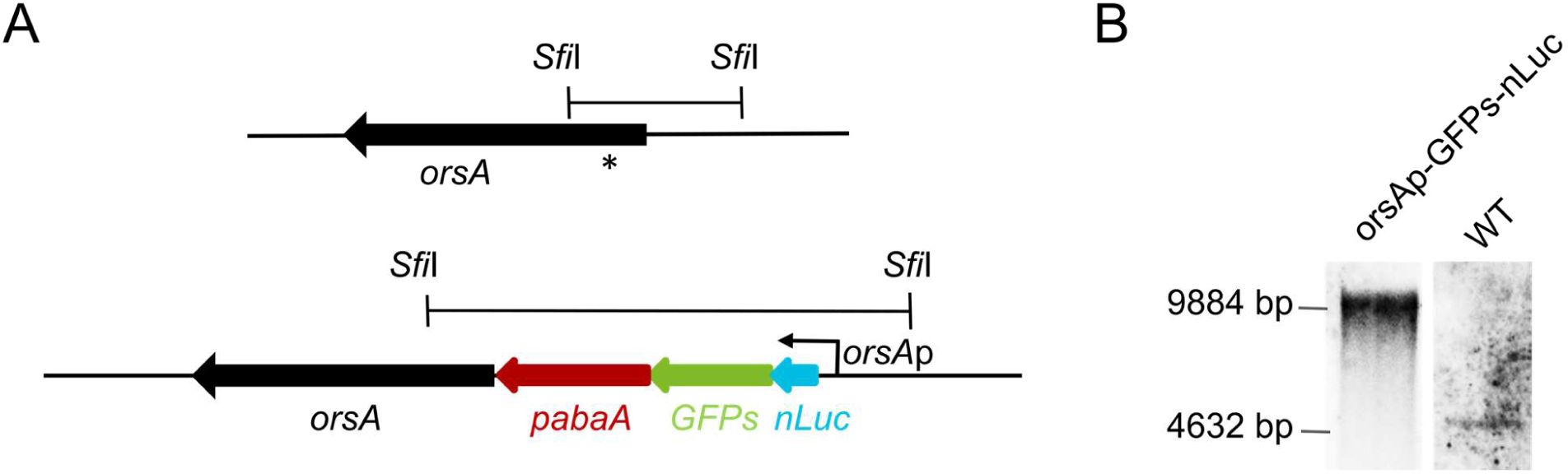
Genomic organization and Southern blot analysis of *A. nidulans orsA*p-nLuc- GFPs reporter strain. **A:** Scheme of the *orsA* genomic locus in the wild type (bottom) and the *orsA* genomic locus (top) in the *A. niduans* reporter strain containing the *orsA* promoter 5’ of the nLuc-GFPs translational gene fusion. The probe used for Southern blot analysis is indicated by *. **B:** Southern blot analysis of genomic DNA of the wild-type and *orsA*p-nLuc-GFPs reporter strain. Genomic DNA was digested with *Sfi*I overnight. The probe (*) was directed against the upstream region of the *orsA* gene (see A). The wild-type strain gives a band of 4632 bp, while for the reporter strain a band of 9884 bp is characteristic of the integrated gene fusion at the *orsA* locus.

**Supp. Fig.7:**
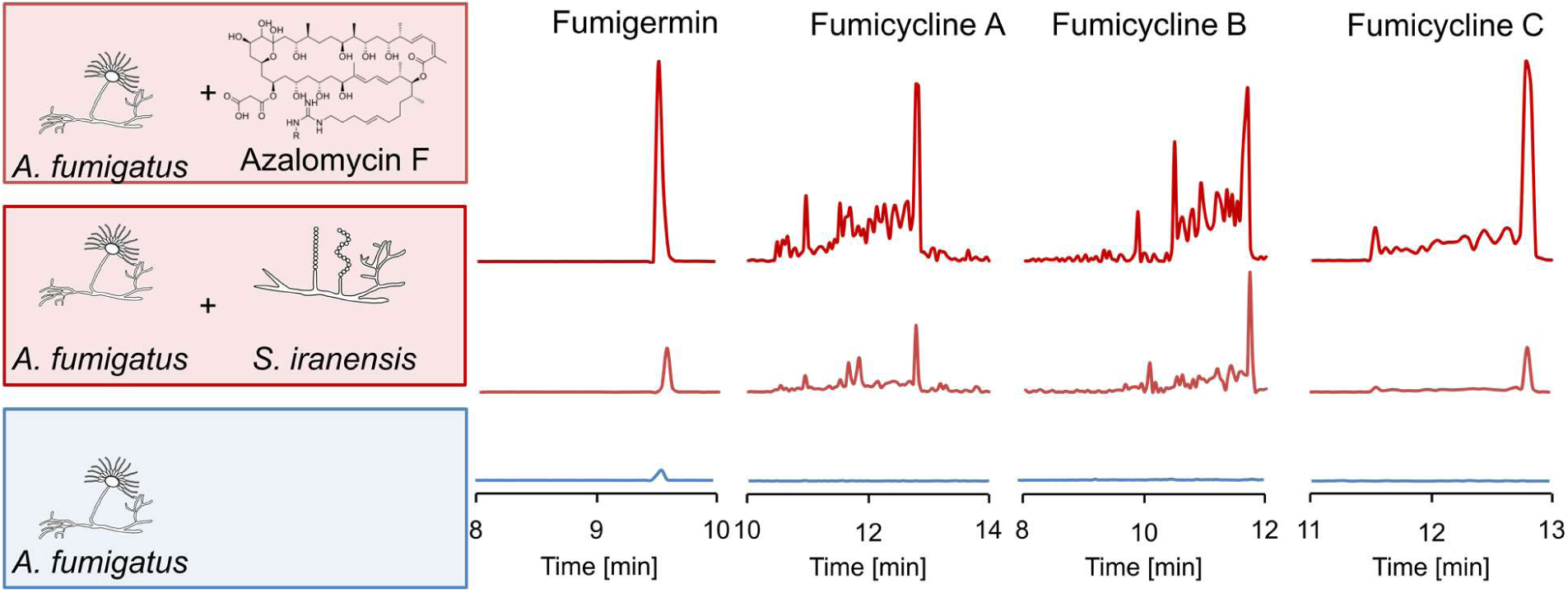
Production of fumigermin and fumicycline A - C in *A. fumigatus* triggered by azalomycin F. Cultures of *A. fumigatus* were supplemented with azalomycin F or, as a control, co-incubated with *S. iranensis*. Extracted ion chromatograms for fumigermin (*m/z* 195 [M+H]^+^), fumicycline A (*/mz* 423 [M-H]^-^), fumicycline B (*/mz* 441 [M-H]^-^) and fumicycline C (*/mz* 483 [M-H]^-^) derived from LC-MS analysis of culture extracts are shown.

**Supp. Fig.8.**
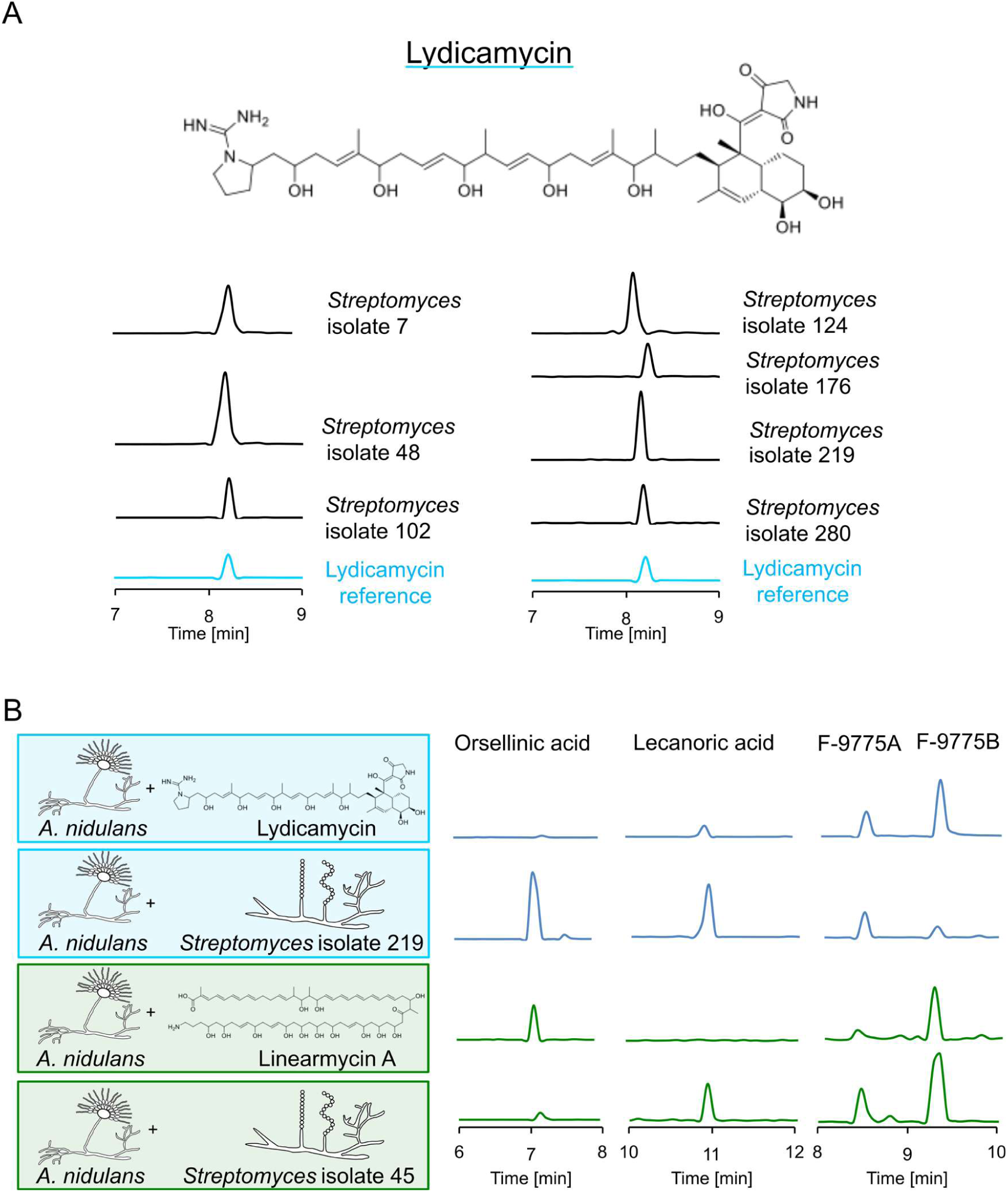
Identification of lydicamycin and linearmycin as well as their producing streptomycetes as inducers of production of orsellinic acid and derivatives in *A. nidulans*. **A:** Cultivation of indicated *Streptoymces* strains isolated from soil and extracted ion chromatogram for lydicamycin (*/mz* 841 [M+H]^+^)) derived from LC-MS analysis of culture extracts compared to standard. The structure of lydicamycin is shown on top. **B:** *A. nidulans* cultures supplemented with indicated compounds or co-incubated with lydicamycin-producing *Streptomyces* isolate 219 and linearmycin A-producing *Streptomyces* isolate 45. Extracted ion chromatograms of orsellinic acid (*m/z* 167 [M-H]^-^), lecanoric acid (*m/z* 317 [M-H]^-^), and compounds F-9775A / F-9775B (*m/z* 395 [M-H]^-^) are shown.

**Supp. Fig.9.**
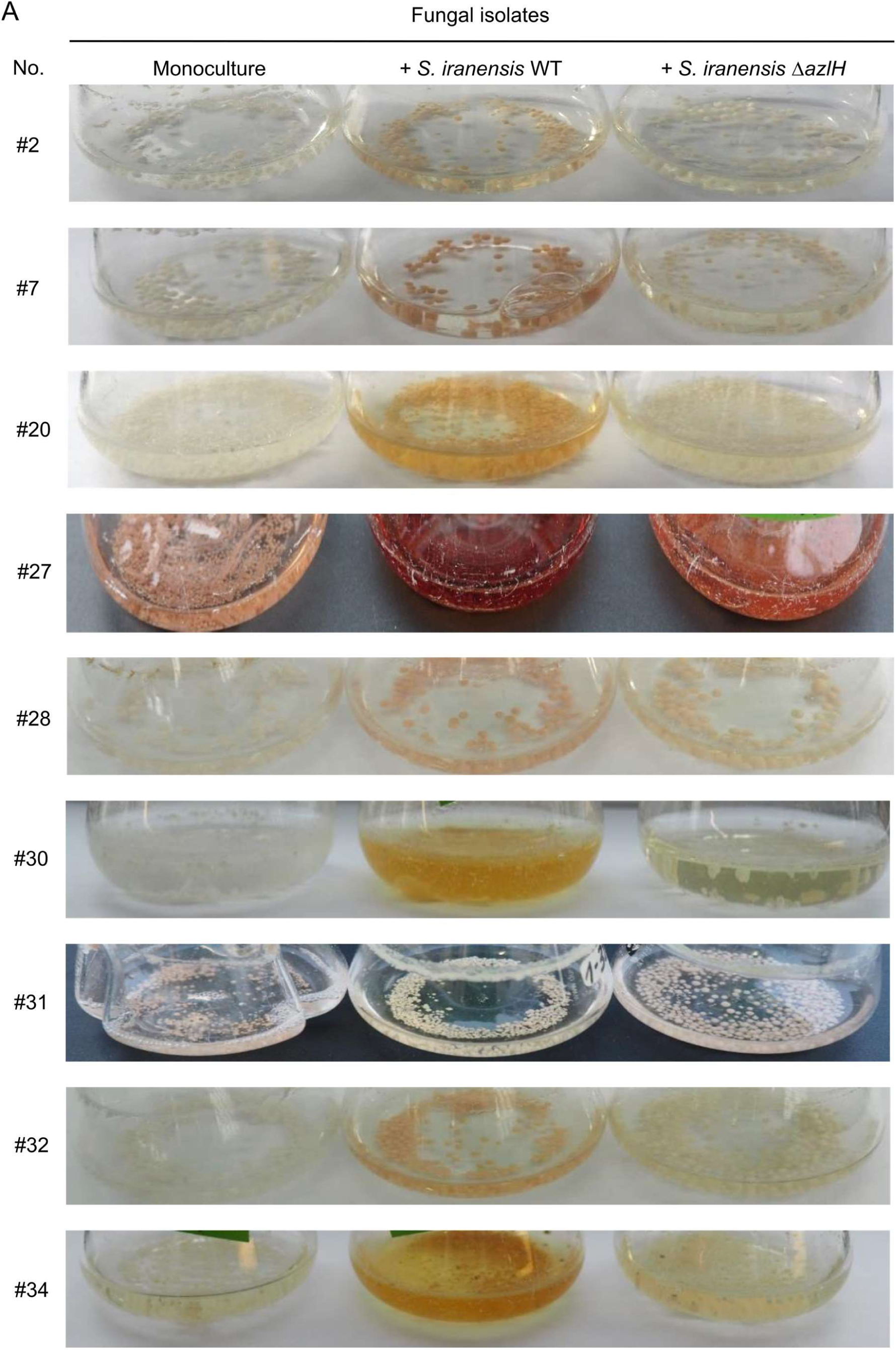
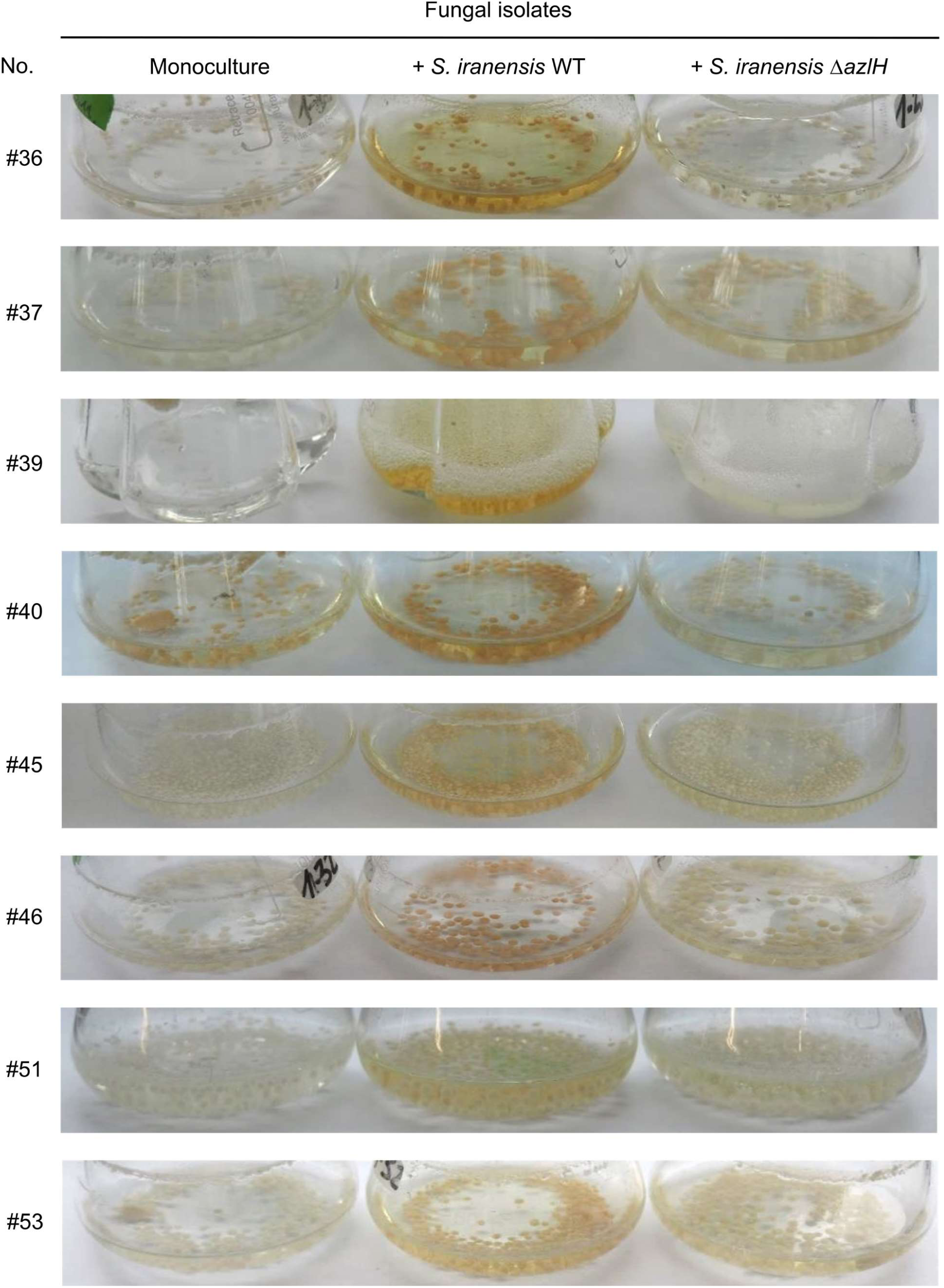
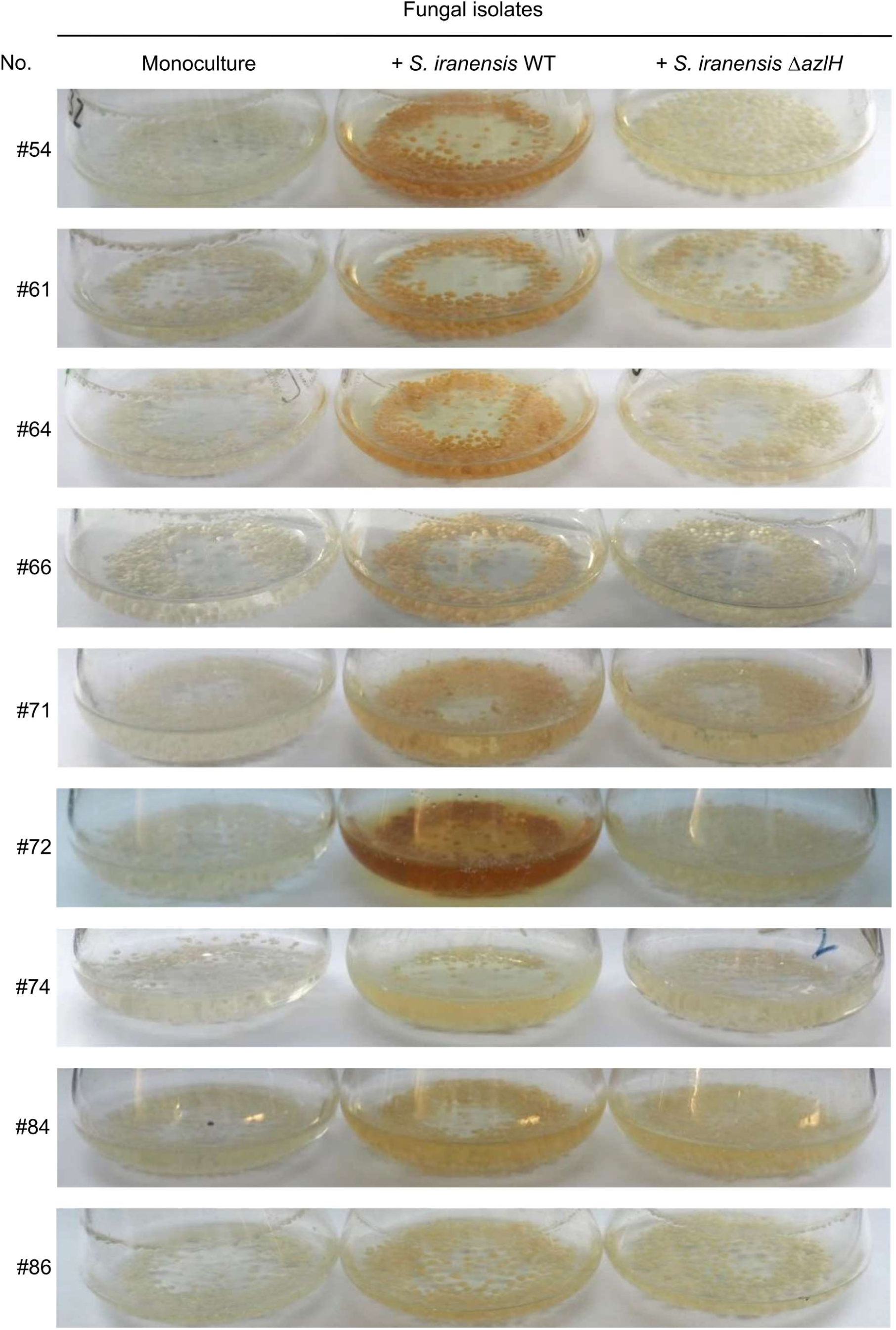
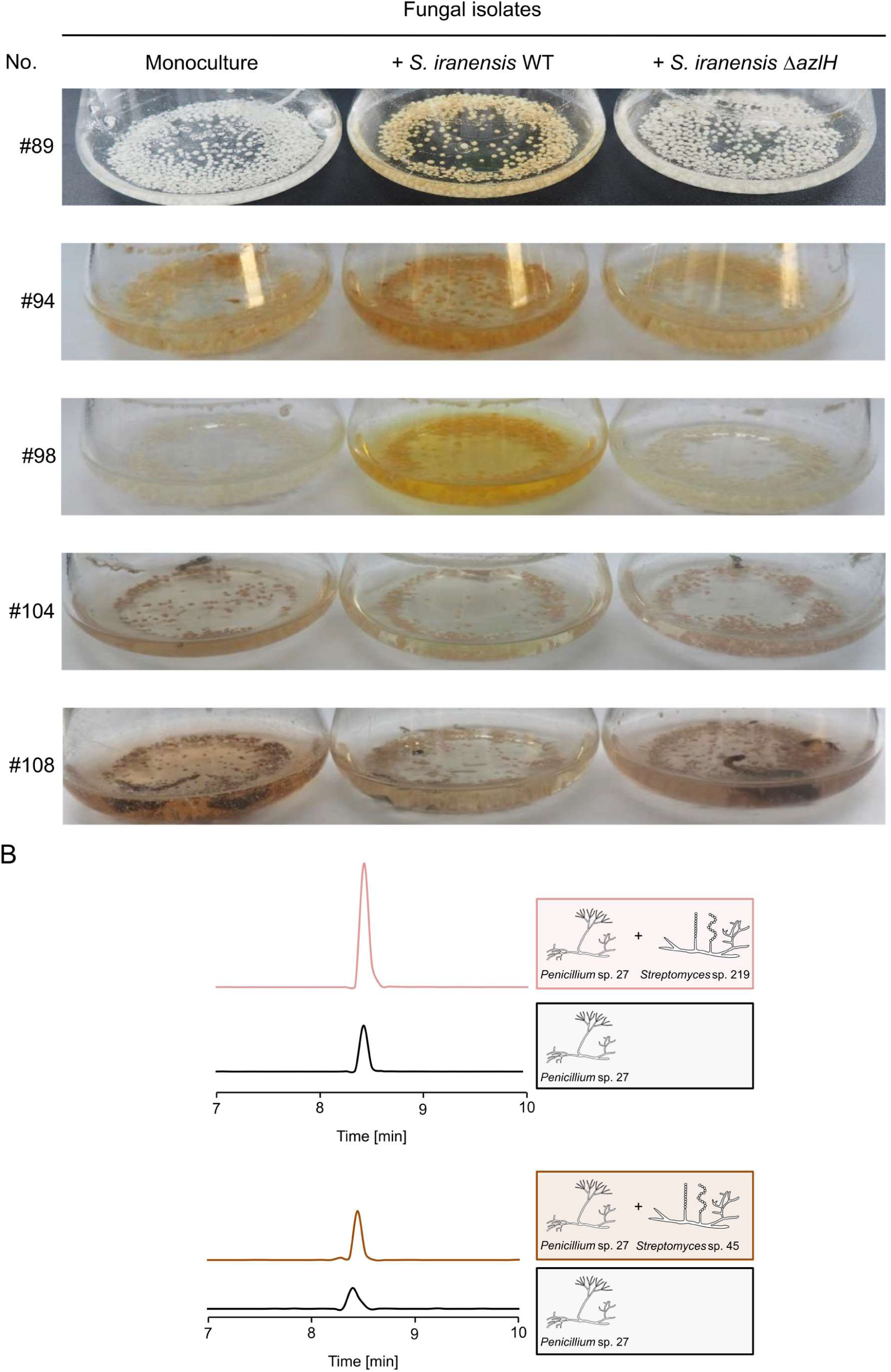
Identification of fungal soil isolates differently responding to *S. iranensis* WT and the azalomycin F-deficient Δ*azlH* mutant strain. **A**: Cultivation flasks with monocultures and cocultures of fungal soil isolates with *S. iranensis* WT and the azalomycin F-deficient mutant strain **Δ***azlH* are shown. **B**: Extracted ion chromatograms for carviolin (*m/z* 299 [M–H]^-^) derived from LC-MS analysis of culture extracts of mono- and co-cultivation of *Penicillium* sp. 27 with *Streptomyces* sp. 45 *and Streptomyces* sp. 219

**Supp. Fig.10.**
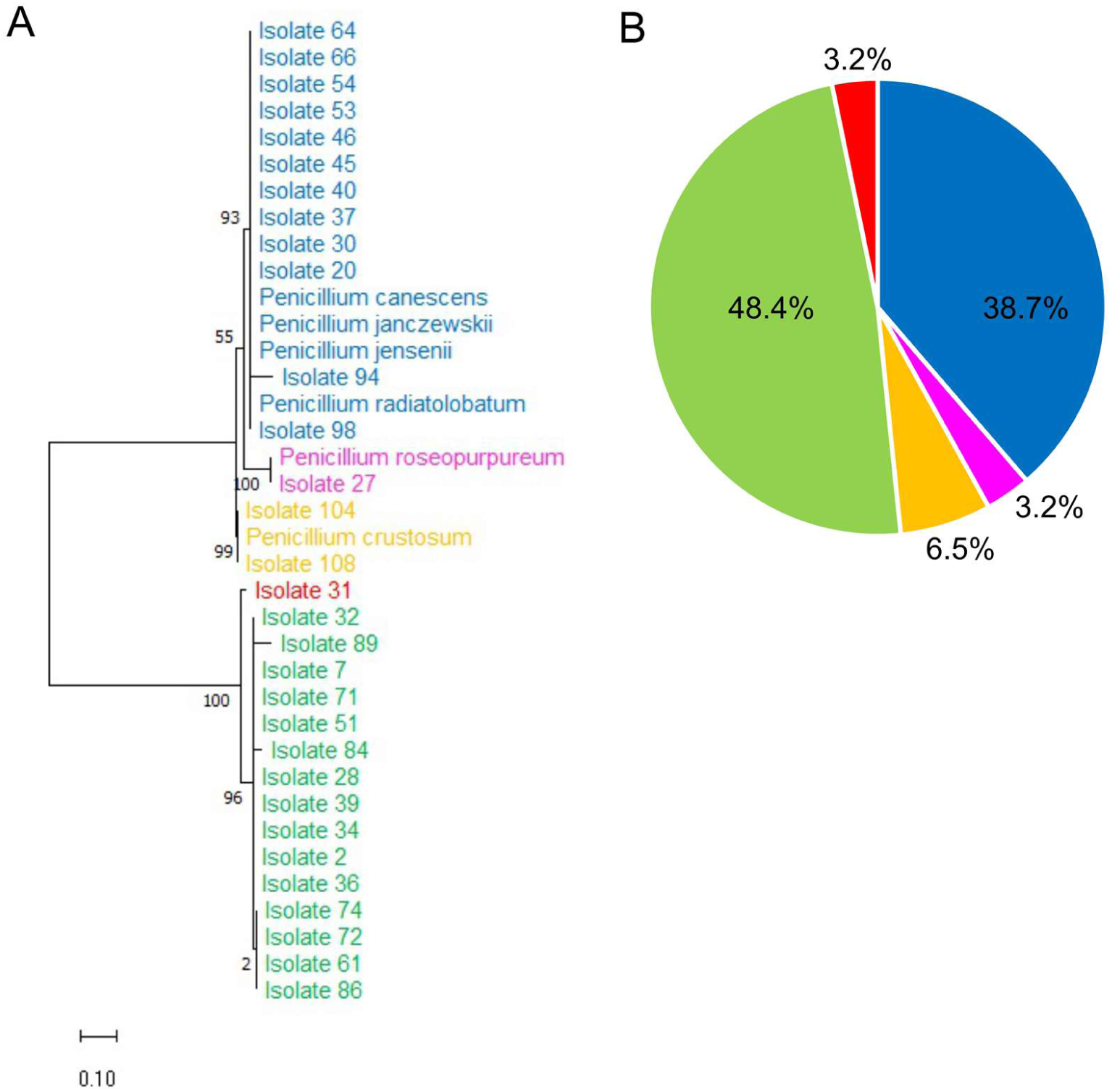
Phylogenetic tree of fungal isolates responding to arginine-derived polyketides. **A:** Phylogenetic tree of fungal isolates based on program MEGA X. The Maximum Likelihood method and the Jukes-Cantor model were used. Bootstrap values obtained after 500 replications are indicated at nodes. The tree is drawn to scale, with branch length measured in the number of base substitutions per site. **B**: Frequency of isolation of fungal species and unknown species from soil. Groups 1 and 2 are defined as “unknown” by, based on their ITS sequences, not grouping with each other and with a type strain from the NCBI database. ITS sequences are will be made accessible for the final publication.

**Supp. Fig.11:**
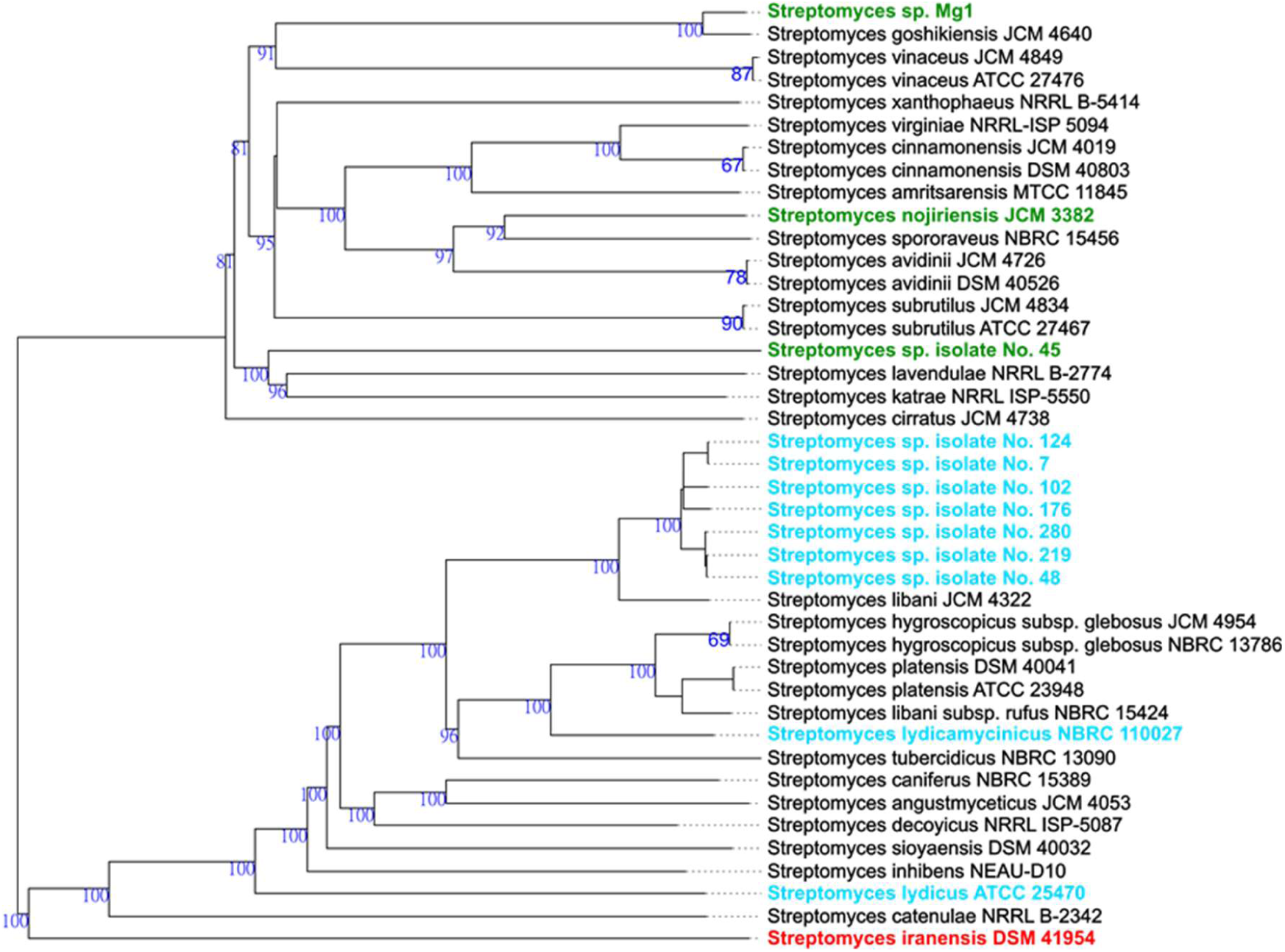
Phylogenetic tree of arginine-derived polyketide producers with a focus on strains isolated here. Genomes of soil isolates producing lydicamycin and linearmycin were compared to known producers of these compounds and *S. iranensis* using the Type Strain Genome Server (TYGS) provided by the DSMZ, Germany (*50*). The genomes of the streptomycetes will be made accessible for the final publication.

**Supp. Fig.12:**
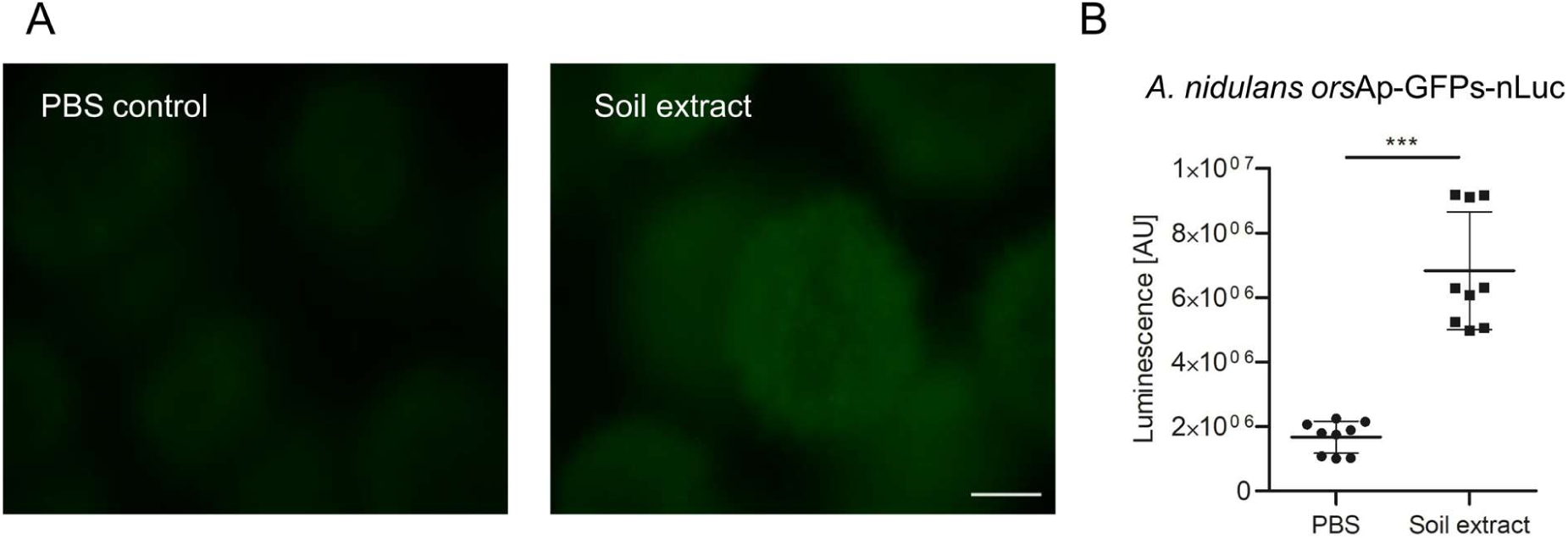
Activation of the *orsA* promoter of *A. nidulans* by soil extract. **A:** Fluorescence images of *A. nidulans orsA*p-nLuc-GFPs reporter strain incubated for 24 h with PBS (PBS control) or soil extract. Scale bar: 500 µm. **B**: Luminescence due to nano-luciferase of *A. nidulans orsA*p-nLuc-GFPs reporter strain after addition of PBS (control) or a PBS extract of soil used here for the isolation of fungi and filamentous bacteria. Error bars represent the mean ± SEM. *** *p* ≤ 0.005 (unpaired, two-tailed t test).

**Supp. Fig.13.**
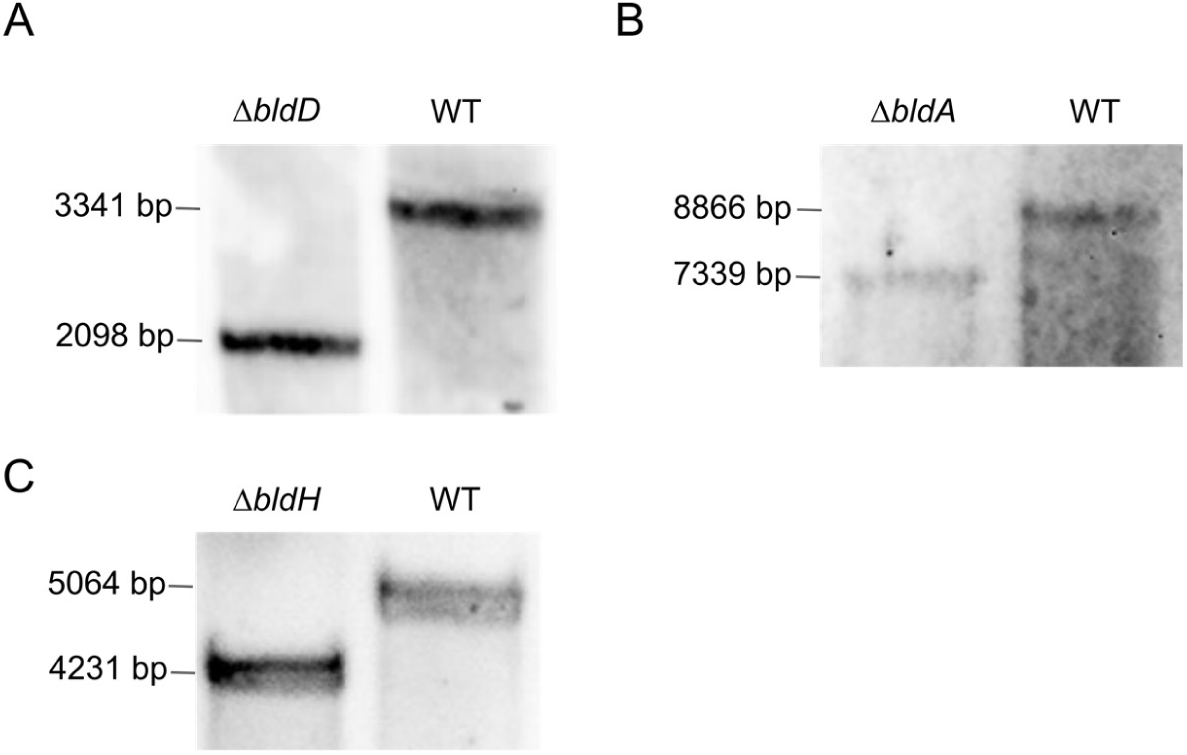
Southern blot analysis of the generated *S. iranensis* deletion mutants. Each blot shows the band characteristic of the indicated deleted gene in comparison to a band obtained with *S. iranensis* wild type. Genomic DNA was digested with *Bam*HI for verification of *ΔbldA*, *Bst*EII for Δ*bldD*, and *Pst*I for *ΔbldH*. **A:** Deletion of *bldD*. The expected band for a deletion reflects a size of 2098 bp, for the wild type 3341 bp. **B:** Deletion of *bldA.* Deletion of *bldA* is indicated by a band of 7339 bp, for the wild type 8866 bp. **C:** Deletion of *bldH.* Insertion of the resistance cassette is reflected by a band reflecting a size of 4231 bp, for the wild type 5064 bp.

## Notes

### Competing Interest Statement

The authors have declared no competing interest.

